# The Landscape-Based Protein Stability Analysis and Network Modeling of Multiple Conformational States of the SARS-CoV-2 Spike D614 Mutant: Conformational Plasticity and Frustration-Driven Allostery as Energetic Drivers of Highly Transmissible Spike Variant

**DOI:** 10.1101/2021.12.09.471953

**Authors:** Gennady Verkhivker, Steve Agajanian, Ryan Kassab, Keerthi Krishnan

**Affiliations:** Keck Center for Science and Engineering, Graduate Program in Computational and Data Sciences, Schmid College of Science and Technology, Chapman University, Orange, CA 92866, United States of America; Department of Biomedical and Pharmaceutical Sciences, Chapman University School of Pharmacy, Irvine, CA 92618, United States of America

## Abstract

The structural and functional studies of the SARS-CoV-2 spike protein variants revealed an important role of the D614G mutation that is shared across many variants of concern(VOCs), suggesting the effect of this mutation on the enhanced virus infectivity and transmissibility. The recent structural and biophysical studies provided important evidence about multiple conformational substates of the D614G spike protein. The development of a plausible mechanistic model which can explain the experimental observations from a more unified thermodynamic perspective is an important objective of the current work. In this study, we employed efficient and accurate coarse-grained simulations of multiple structural substates of the D614G spike trimers together with the ensemble-based mutational frustration analysis to characterize the dynamics signatures of the conformational landscapes. By combining the local frustration profiling of the conformational states with residue-based mutational scanning of protein stability and network analysis of allosteric interactions and communications, we determine the patterns of mutational sensitivity in the functional regions and sites of variants. We found that the D614G mutation may induce a considerable conformational adaptability of the open states in the SARS-CoV-2 spike protein without compromising folding stability and integrity of the spike protein. The results suggest that the D614G mutant may employ a hinge-shift mechanism in which the dynamic couplings between the site of mutation and the inter-protomer hinge modulate the inter-domain interactions, global mobility change and the increased stability of the open form. This study proposes that mutation-induced modulation of the conformational flexibility and energetic frustration at the inter-protomer interfaces may serve as an efficient mechanism for allosteric regulation of the SARS-CoV-2 spike proteins.

## Introduction

SARS-CoV-2 infection is transmitted when the viral spike (S) glycoprotein binds to the host cell receptor ACE2, leading to the entry of S protein into host cells and membrane fusion.^1, 2^ The full-length SARS-CoV-2 S protein consists of amino (N)-terminal S1 subunit and carboxyl (C)-terminal S2 subunit where S1 is involved in the interactions with the host receptor and includes an N-terminal domain (NTD), the receptor-binding domain (RBD), and two structurally conserved subdomains (SD1 and SD2). The S2 subunit is highly evolutionary conserved and contains an N-terminal hydrophobic fusion peptide (FP), fusion peptide proximal region (FPPR), heptad repeat 1 (HR1), central helix region (CH), connector domain (CD), heptad repeat 2 (HR2), transmembrane domain (TM) and cytoplasmic tail (CT) regions (Figure S1). Structural and biochemical studies established that the mechanism of virus infection may involve conformational transitions between distinct functional forms of the SARS-CoV-2 S protein in which the RBDs continuously switch between “down” and “up” positions.^3–10^ The cryo-EM structure of the SARS-CoV-2 S trimer revealed a spectrum of closed states that included a structurally rigid closed form and more dynamic closed states preceding a transition to the fully open S conformation.^5^ Protein engineering and structural studies also showed that specific disulfide bonds and proline mutations can modulate stability of the SARS-CoV-2 S trimer^6^ and lead to the thermodynamic shifts between the closed and open forms.^7–9^ Dynamic structural changes that accompany SARS-CoV-2 S binding with the ACE2 host receptor were described in cryo-EM experiments showing a cascade of conformational transitions from a compact closed form weakened after furin cleavage to the partially open states and subsequently the ACE2-bound open form thus priming the S protein for fusion.^10^ The cryo-EM and tomography examined conformational flexibility and distribution of S trimers in situ on the virion surface^11^ showing that spontaneous conformational changes and population shifts between different functional states can be maintained in different biological environments, reflecting the intrinsic properties of conformational landscapes for SARS-CoV-2 S trimers. Single-molecule Fluorescence (Förster) Resonance Energy Transfer (smFRET) studies of SARS-CoV-2 S trimer on virus particles revealed a sequence of conformational transitions from the closed state to the receptor-bound open state suggesting that mechanisms of conformational selection and receptor-induced structural adaptation can work synchronously and showing that SARS-CoV-2 neutralization can be achieved by antibodies that either directly compete with the ACE2 receptor binding or exert their effect by allosterically stabilizing the S protein in its RBD-down conformation.^12^

The vast and continuously expanding body of structural and biochemical studies of the SARS-CoV-2 S complexes with different classes of potent antibodies targeting distinct binding epitopes of the S-RBD as well as various antibody cocktails and combinations have revealed multiple conformation-dependent epitopes, highlighting the link between conformational plasticity and adaptability of S proteins and capacity for eliciting specific binding and broad neutralization responses.^13–36^ Deep mutagenesis scanning of antibody-escaping mutations showed that the escape mutations cluster in several RBD regions and can be constrained with respect to their effects on expression of properly folded RBD and ACE2 binding.^37–42^ Functional and structural studies explored how B1.1.7 (alpha), B.1.351 (beta), P1 (gamma) and B1.1.427/B.1.429 (epsilon) variants in the SARS-CoV-2 S protein affect conformational landscapes of the S protein and the ability to evade host immunity and incur resistance to antibodies.^43–53^ These studies revealed subtle structural and functional impact of mutations that can modulate dynamics and stability of the closed and open forms, increase binding to the human receptor ACE2, and confer resistance to neutralizing antibodies.

Strikingly, all these emerging variants of SARS-CoV-2 designated into variants of concern(VOCs) share D614G mutation, that individually was linked with the enhanced infectivity profile and attracted significant early attention following the evidence of the mutation enrichment via epidemiological surveillance.^54, 55^ The initial structural studies showed that the D614G mutation can act by shifting the population of the SARS-CoV-2 S trimer from the closed form (53% of the equilibrium) in the native spike protein to a widely-open topology of the “up” protomers in the D614G mutant with 36% of the population adopting a single open protomer, 39% with two open protomers and 20% with all three protomers in the open conformation.^56^ The cryo-EM structures of the S-D614 and S-G614 ectodomains showed the increased population of the 1-RBD-up open form as compared to the closed state in the S-GSAS/D614G structure.^57^ The electron microscopy analysis also revealed the higher 84% percentage of the 1-up RBD conformation in the S-G614 protein.^58^ Functional studied showed that the S-G614 mutant exhibited the greater infectivity than the S-D614 protein which was attributed to the greater stability of the S-G614 mutant and leading to the reduced S1 subdomain shedding.^59^ An alternative mechanistic scenario proposed that D614G mutation would promote rather than limit shedding of the S1 domain by altering the transitions of “up” and “down” forms.^60^ The increased stability of the D614G mutant was inferred from the recent cryo-EM structures of a full-length unmodified S-G614 trimer that can reversibly adopt an all-down closed state, an intermediate (1 RBD in an intermediate state), and 1 RBD-up open conformation.^61^ These studies showed that the S-D614 protein was less stable, while the S-G614 mutant was stabilized in the 1 RBD-up conformations that featured ordering of the so-called 630-loop (residues 620-640) near the interface of S1 and S2 thus helping to strengthen the inter-domain interactions and enhance the stability of the mutated S protein. This study provided support to the reduced shedding mechanism induced by the D614G mutation that inhibits a premature dissociation of the S1 subunit which eventually leads to the increased number of functional spikes and stronger infectivity.^61^ Structure-based protein design and cryo-EM structure determination provided an additional support to the stability hypothesis showing that D614G and D614N mutations can result in the increased stability of the S protein mutants and decrease the premature shedding of the S1 domain.^62^ The recent series of cryo-EM studies have further expanded structural characterization of the S-G614 trimer conformations, revealing a range of more flexible closed and open conformations, showing that the S-G614 occupies predominantly the open conformation with 87 % of the population in the 1 RBD-up or 2-RBD-up forms compared with 17% in the D614 structure for which 2 RBD-up form was never detected.^63^ Cryo-EM structures of trimeric S protein in multiple states including the all RBD-locked, activated, S1/S2 trypsin-cleaved, and the full-length ACE2-bound states showed that S-G614 and the trypsin-cleaved S-G614 proteins adopt 1 RBD-up conformation confirming that D614G mutation can increase the flexibility and population of the open spike states which might contribute to the increased infectivity.^64^ By combining cryo-EM analysis with differential scanning calorimetry and differential scanning fluorimetry, the recent illuminating investigation characterized multiple substates of the of the S-G614 protein in open and closed forms, showing a significant conformational heterogeneity and thermodynamic preferences of the S-G614 for 1 RBD-up and 2 RBD-up open states.^65^ Using an array of biophysical tools, this study convincingly demonstrated the increased stability of the S-G614 trimer manifested in the increased resistance to cold-and heat-induced unfolding, and suggested that the increased conformational flexibility and emergence of multiple substates of the open form may be the main thermodynamic driver promoting virus infectivity.^65^ These findings were echoed in another biophysical study showing that the S-G614 variant was more stable than the S-D614 variant and had better binding ability with the ACE2 receptor after storage at -20°C for up to 30 days.^66^ This study supported the notion that the increased stability and infectivity of the S-G614 variant are intimately connected and the higher stability of S-G614 compared to that of S-D614 may contribute to rapid viral spread.^66^ The biophysical experiments using biolayer interferometry technology revealed that the S-G614 variant displays highest entry efficiency among natural S variants and the increased binding affinity for the ACE2 protein dimer as compared to S-D614 protein in a temperature-dependent manner, suggesting that mutation-induced conformational flexibility of the S protein may be responsible for the observed ∼2.0 -fold increase in the binding affinity.^67^

The multiscale structural modeling and atomistic molecular dynamics (MD) simulations of the full-length SARS-CoV-2 S glycoprotein embedded in the viral membrane, with a complete glycosylation profile provided the blueprint for biophysical modeling of S proteins.^68–74^ Using deep data analysis and protein structure network modeling of MD simulations, another computational study identified residues that exhibit long-distance coupling with the RBD opening, including sites harboring functional mutations D614G and A570D which points to the important role of D614G variant in modulating allosteric communications in the S protein.^75^ The free energy landscapes of the S protein derived from MD simulations together with nudged elastic pathway optimization mapping of the RBD opening revealed a specific transient allosteric pocket at the hinge region that is located near D614 position influences RBD dynamics.^76^ Using computational modeling it was suggested that the D614G mutation may affect the inter-protomer energetics and induce specific interaction changes in the C-terminal domain 1(CTD1) (residues 528-591) and fusion peptide (FP) regions that mediate couplings between S1 and S2 subunits.^77^ MD simulations probed the effects of the D614G mutation, suggesting that the variant favors an open conformation in which S-G614 protein maintains the symmetry in the number of persistent contacts between the three protomers.^78^ Markov modeling characterized the dynamics of the S protein and mutational variants, predicting the increase in the open state occupancy for the D614G mutation due to the increased flexibility of the closed state and the enhanced stabilization of the open form. ^79^ The energy analysis of the S-D614 and S-G614 proteins in the closed and partially open conformations showed that local interactions near D614 position may be energetically frustrated and become stabilized in the S-G614 mutant through strengthening of the inter-protomer association between S1 and S2 regions.^80^ Structure-based model showed that the D614G mutation may promote closer association and stronger interactions with S2 subunit, supporting the reduced shedding mechanism of the increased S-614 stability.^81^ Molecular simulations and network modeling approaches were used on our most recent investigation to present evidence that the SARS-CoV-2 spike protein can function as an allosteric regulatory engine that fluctuates between dynamically distinct functional states.^82, 83^ Allosteric models of the SARS-CoV-2 S protein dynamics proposed in our studies have allowed for better understanding distant couplings between S protein regions and detection of allosteric hotspot switches regulating binding with antibodies and nanobodies.^84–89^

The existing mechanisms suggest that mutation-induced allosteric transitions to the open states and the reduced shedding mechanism of the thermodynamic stability of the open S-G614 trimers are important driving forces for greater infectivity of the D614G mutant. However, a consensus view on the mechanism underlying the functional effects and increased infectivity of S-D614G spike mutant is yet to be established. The development of a plausible mechanistic model which can explain the experimental observations from a unified dynamic perspective that considers local effects and long-range allosteric interactions is an important objective of current investigations. In this study, we employed the efficient and accurate coarse-grained simulations of multiple structural substates of the S-G614 trimers to characterize the dynamics signatures of the conformational landscapes of spike proteins. By combining atomistic simulations with the ensemble-based local frustration analysis of the conformational states, we determine sensitivity of functional regions and sites of mutational variants to energetic variations in different functional states. Using the ensemble-based local frustration profiling and mutational scanning of protein stability changes in the S-G614 conformational states, we quantify conformational plasticity and adaptability of the S-G614 states, showing that the inter-protomer interactions can be generally weakened in significantly heterogeneous and expanded S-G614 open states. Combined, the ensemble-based dynamics and energetic analysis of multiple S-G614 substates suggests a significant conformational and mutational plasticity of the open S-G614 trimer. We show that disorder-order changes in the flexible loop near G614 may be coupled with the energetic frustration in the mutational site and allow for modulation of the inter-protomer interactions. This study suggests that modulation of the energetic frustration at the inter-protomer interfaces by D614G mutation can serve as a mechanism for allosteric regulation in which the dynamic couplings between the site of mutation and the inter-protomer hinge of functional motions would modulate the inter-domain interactions, global changes in mobility and the increased stability of the open form.

## Materials and Methods

### Structure Preparation and Analysis

All structures were obtained from the Protein Data Bank.^90, 91^ During structure preparation stage, protein residues in the crystal structures were inspected for missing residues and protons. Hydrogen atoms and missing residues were initially added and assigned according to the WHATIF program web interface.^92, 93^ The structures were further pre-processed through the Protein Preparation Wizard (Schrödinger, LLC, New York, NY) and included the check of bond order, assignment and adjustment of ionization states, formation of disulphide bonds, removal of crystallographic water molecules and co-factors, capping of the termini, assignment of partial charges, and addition of possible missing atoms and side chains that were not assigned in the initial processing with the WHATIF program. The missing loops in the studied cryo-EM structures of the SARS-CoV-2 S protein were reconstructed and optimized using template-based loop prediction approaches ModLoop^94^, ArchPRED Server^95^ and further confirmed by FALC (Fragment Assembly and Loop Closure) program.^96^ The side chain rotamers were refined and optimized by SCWRL4 tool.^97^ The conformational ensembles were also subjected to all-atom reconstruction using PULCHRA method^98^ and CG2AA tool^99^ to produce atomistic models of simulation trajectories. The protein structures were then optimized using atomic-level energy minimization with a composite physics and knowledge-based force fields as implemented in the 3Drefine method.^100^ The atomistic structures from simulation trajectories were further elaborated by adding N-acetyl glycosamine (NAG) glycan residues and optimized. The glycosylated microenvironment for atomistic models of the simulation trajectories was mimicked by using the structurally resolved glycan conformations for 22 most occupied N-glycans^101, 102^ in each as determined in the cryo-EM structures of the SARS-CoV-2 spike S trimer in the closed state (K986P/V987P,) (pdb id 6VXX) and open state (pdb id 6VYB), and the cryo-EM structure SARS-CoV-2 spike trimer (K986P/V987P) in the open state (pdb id 6VSB).

### Coarse-Grained Simulations and Elastic Models

Coarse-grained (CG) models are computationally effective approaches for simulations of large systems over long timescales. We employed CABS-flex approach that efficiently combines a high-resolution coarse-grained model and efficient search protocol capable of accurately reproducing all-atom MD simulation trajectories and dynamic profiles of large biomolecules on a long time scale.^103–109^ In the CG-CABS model, the amino acid residues are represented by Cα, Cβ, the center of mass of side chains and another pseudoatom placed in the center of the Cα-Cα pseudo-bond. The sampling scheme of the CABS model used in our study is based on Monte Carlo replica-exchange dynamics and involves a sequence of local moves of individual amino acids in the protein structure as well as moves of small fragments.^103–105^ CABS-flex standalone package dynamics implemented as a Python 2.7 object-oriented package was used for fast simulations of protein structures.^107–109^ In the CABS-flex package we also used MODELLER-based reconstruction of generated models and simulation trajectories to all-atom representation. The default settings were applied in which soft native-like restraints are imposed only on pairs of residues fulfilling the following conditions : the distance between their *C*^α^ atoms was smaller than 8 Å, and both residues belong to the same secondary structure elements. A total of 1,000 independent CG-CABS simulations were performed for each of the studied systems. In each simulation, the total number of cycles was set to 10,000 and the number of cycles between trajectory frames was 100.

We performed principal component analysis (PCA) of reconstructed trajectories derived from CABS-CG simulations using the CARMA package^110^ and also determined the essential dynamics profiles in slow modes using elastic network models (ENM) analysis.^111^ Two elastic network models: Gaussian network model (GNM)^112, 113^ and Anisotropic network model (ANM) approaches^114^ were adopted to compute the amplitudes of isotropic thermal motions and directionality of anisotropic motions. The functional dynamics analysis was conducted using the GNM in which protein structure is reduced to a network of *N* residue nodes identified by *C_α_* atoms and the fluctuations of each node are assumed to be isotropic and Gaussian. The topology of the protein structure is described by *N*×*N* Kirchhoff matrix of inter-residue contacts Г, where the off-diagonal elements are −1, if the nodes are within a cutoff distance *r_c_*, and zero otherwise. In GNM approach the interaction potential for a protein of *N* residues was:

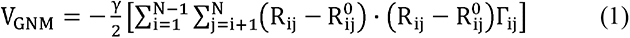

where 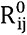 and R_ij_ were the equilibrium and instantaneous distance between residues i and j, and *Г* was the N×N Kirchhoff matrix:

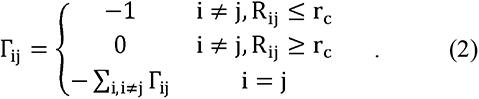

Thus, the square fluctuations were:

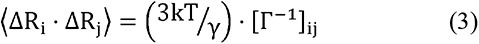

The normal modes were extracted by eigenvalue decomposition of the matrix *Г* = *U*Λ*U^T^*, *U* being the orthogonal matrix whose k^th^ column u_k_ was the k^th^ mode eigenvector. Λ was the diagonal matrix of eigenvalues λ_k_.

In ANM approach^92^ the interaction potential for a protein of *N* residues was:

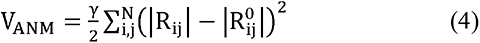

The 3*N*×3*N* Hessian matrix, *H*, determines the systems dynamics.

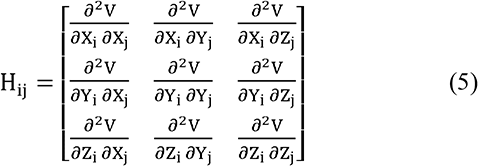

where *X_i_, Y_i_* and *Z_i_* represented the Cartesian components of the i-th residue, *V* was the potential energy of the system. We selected r_c_ =13 Å. ANMs provide information about the amplitudes, as well as the direction of residue fluctuations. Conformational mobility profiles in the essential space of low frequency modes were obtained using DynOmics server^113^ and ANM server.^114^

### Mutational Sensitivity Analysis and Alanine Scanning

To compute protein stability changes in the SARS-CoV-2 S structures, we conducted a systematic alanine scanning of protein residues in the SARS-CoV-2 trimer mutants as well as mutational sensitivity analysis at the mutational site for both SARS-CoV-2 S-D614 and SARS-CoV-2 S-G614 structures. Two different approaches were used. Alanine scanning of protein residues was performed using FoldX approach.^115–120^ and BeAtMuSiC approach.^121–123^ If a free energy change between a mutant and the wild type (WT) proteins ΔΔG= ΔG (MT)-ΔG (WT) > 0, the mutation is destabilizing, while when ΔΔG <0 the respective mutation is stabilizing. BeAtMuSiC approach is based on statistical potentials describing the pairwise inter-residue distances, backbone torsion angles and solvent accessibilities, and considers the effect of the mutation on the strength of the interactions at the interface and on the overall stability of the complex.^121–123^ We leveraged rapid calculations based on statistical potentials to compute the ensemble-averaged alanine scanning computations and mutational sensitivity analysis at D614 and G614 positions using equilibrium samples from reconstructed simulation trajectories.

### Dynamic-Based Modeling of Residue Interaction Network and Community Analysis

A graph-based representation of protein structures^124, 125^ is used to represent residues as network nodes and the inter-residue edges to describe non-covalent residue interactions. The network edges that define residue connectivity are based on non-covalent interactions between residue side-chains that define the interaction strength *I_ij_* according to the following expression used in the original studies.^124, 125^

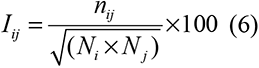

where *n_ij_* is number of distinct atom pairs between the side chains of amino acid residues *i* and *j* that lie within a distance of 4.5 Å. *N_i_* and *N _j_* are the normalization factors for residues *i* and *j* . We constructed the residue interaction networks using both dynamic correlations^126^ and coevolutionary residue couplings^127^ that yield robust network signatures of long-range couplings and communications. The details of this model were described in our previous studies.^128, 129^ In brief, the edges in the residue interaction network are then weighted based on dynamic residue correlations and coevolutionary couplings measured by the mutual information scores.

The edge lengths in the network are obtained using the generalized correlation coefficients *R_MI_*(*X_i_*,*Xj*) associated with the dynamic correlation and mutual information shared by each pair of residues. The length (i.e. weight) *w_ij_* = –log[*R_MI_*(*X_i_*,*X_j_*)] of the edge that connects nodes *i* and *j* is defined as the element of a matrix measuring the generalized correlation coefficient *R_MI_*(*X_i_*,*X_j_*) as between residue fluctuations in structural and coevolutionary dimensions. Network edges were weighted for residue pairs with *R_MI_*(*X_i_*,*X_j_*) > 0.5 in at least one independent simulation as was described in our initial study. The matrix of communication distances is obtained using generalized correlation between composite variables describing both dynamic positions of residues and coevolutionary mutual information between residues. As a result, the weighted graph model defines a residue interaction network that favors a global flow of information through edges between residues associated with dynamics correlations and coevolutionary dependencies.

To characterize allosteric couplings of the protein residues and account for cumulative effect of dynamic and coevolutionary correlations, we employed the generalized correlation coefficient first proposed by Lange and Grubmüller.^130^ The g_correlation tool in the Gromacs 3.3 package was used that allows computation of both linear or non-linear generalized correlation coefficients.^131^ This approach has also been utilized in a similar context of allosteric modeling by other groups^132–135^ where the generalized correlation GC matrix measured dynamic interdependencies between on fluctuations of spatially separated residues.

The RING program^136, 137^ was also employed for the initial generation of residue interaction networks. The ensemble of shortest paths is determined from matrix of communication distances by the Floyd-Warshall algorithm.^138^ Network graph calculations were performed using the python package NetworkX.^139^

The Girvan-Newman algorithm^140–142^ is used to identify local communities. In this approach, edge centrality (also termed as edge betweenness) is defined as the ratio of all the shortest paths passing through a particular edge to the total number of shortest paths in the network. The method employs an iterative elimination of edges with the highest number of the shortest paths that go through them. By eliminating edges, the network breaks down into smaller communities. The algorithm starts with one vertex, calculates edge weights for paths going through that vertex, and then repeats it for every vertex in the graph and sums the weights for every edge. However, in complex and dynamic protein structure networks it is often that number of edges could have the same highest edge betweenness.

An improvement of Girvan-Newman method was implemented, and the algorithmic details of this modified scheme were given in our recent studies.^143–145^ Briefly, in this modification of Girvan-Newman method, instead of a single highest edge betweenness removal, all highest betweenness edges are removed at each step of the protocol. This modification makes community structure determination invariant to the labeling of the nodes in the graph and leads to a more stable solution. The modified algorithm proceeds through the following steps : a) Calculate edge betweenness for every edge in the graph; b) Remove all edges with highest edge betweenness within a given threshold; c) Recalculate edge betweenness for remaining edges; d) Repeat steps b-d until graph is empty.

The betweenness of residue *i* is defined as the sum of the fraction of shortest paths between all pairs of residues that pass through residue *i* :

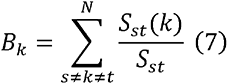

where *S_st_* denotes the number of shortest geodesics paths connecting s and *t,* and *S_st_*(*k*) is the number of shortest paths between residues *s* and *t* passing through the node *k*.

The following Z-score is then calculated

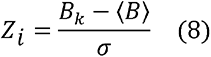

Through mutation-based perturbations of protein residues we compute changes in the average short path length (ASPL) averaged over all possible modifications in a given position. The change of ASPL upon mutational changes of each node is given as

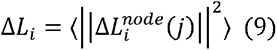

where *i* is a given site, *j* is a mutation and <⋯> denotes averaging over mutations. 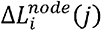 describes the change of ASPL upon mutation *j* in a residue node *i*. Δ*L*_i_ is the average change of ASPL triggered by mutational profiling of this position.

Z-score is then calculated for each node as follows:

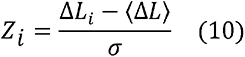

The ensemble-averaged Z –scores ASPL changes are computed from network analysis of the conformational ensembles using 1,000 snapshots of the simulation trajectory for the native protein system.

## Results and Discussion

### Conformational Mobility Profiles of the SARS-CoV-2 S-G614 Proteins: The D614G Mutation Induces Differential Conformational Plasticity in Multiple Conformational Substates

We employed multiple CG simulations followed by atomistic reconstruction and refinement to provide a comparative analysis of the dynamic landscapes characteristic of the major functional states of the SARS-CoV-2 S trimer. While all-atom MD simulations with the explicit inclusion of the glycosylation shield could provide a rigorous assessment of conformational landscape of the SARS-CoV-2 S proteins, such direct simulations remain to be technically challenging due to the size of a complete SARS-CoV-2 S system embedded onto the membrane. To partially address these computational challenges, and enable analysis of multiple conformational states, we combined CG simulations with atomistic reconstruction and additional optimization by adding the glycosylated microenvironment as described in Materials and Methods. The presented analysis characterized the intrinsic dynamics of the SARS-CoV-2 S proteins in the D614G mutant states, particularly probing the diversity of the open S-D614G conformations (Figures 1, 2).

**Figure 1.**
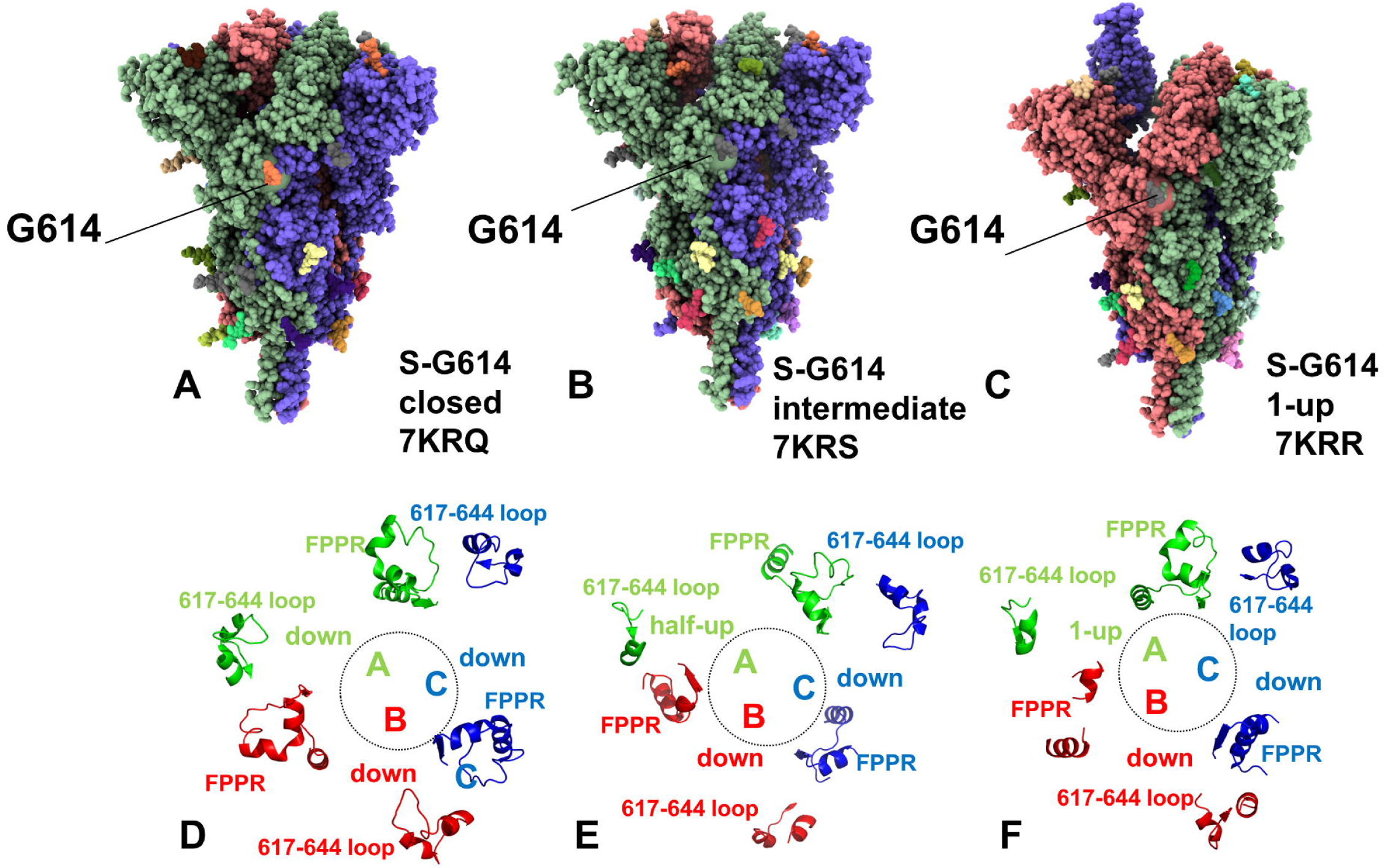
Cryo-EM structures of the SARS-CoV-2 S-G614 trimer structures and composition of the used in this study. The cryo-EM structures of SARS-CoV-2 S-G614 in the closed state, pdb id 7KRQ (A), in the intermediate state, pdb id 7KRS (B) and 1 RBD-up open form, pdb id 7KRR (C). The structures are shown in full spheres and colored with protomers A,B,C are colored in green, red and blue. The position of G614 is shown in spheres. (D-F) Structures of the 630 loop (residues 617 to 644) and FPPR (residues 823 to 862) are shown for protomer A in green ribbons, protomer B in red ribbons and protomer C in blue ribbons. The rendering of SARS-CoV-2 S structures was done using the interactive visualization program UCSF ChimeraX package.^146^

**Figure 2.**
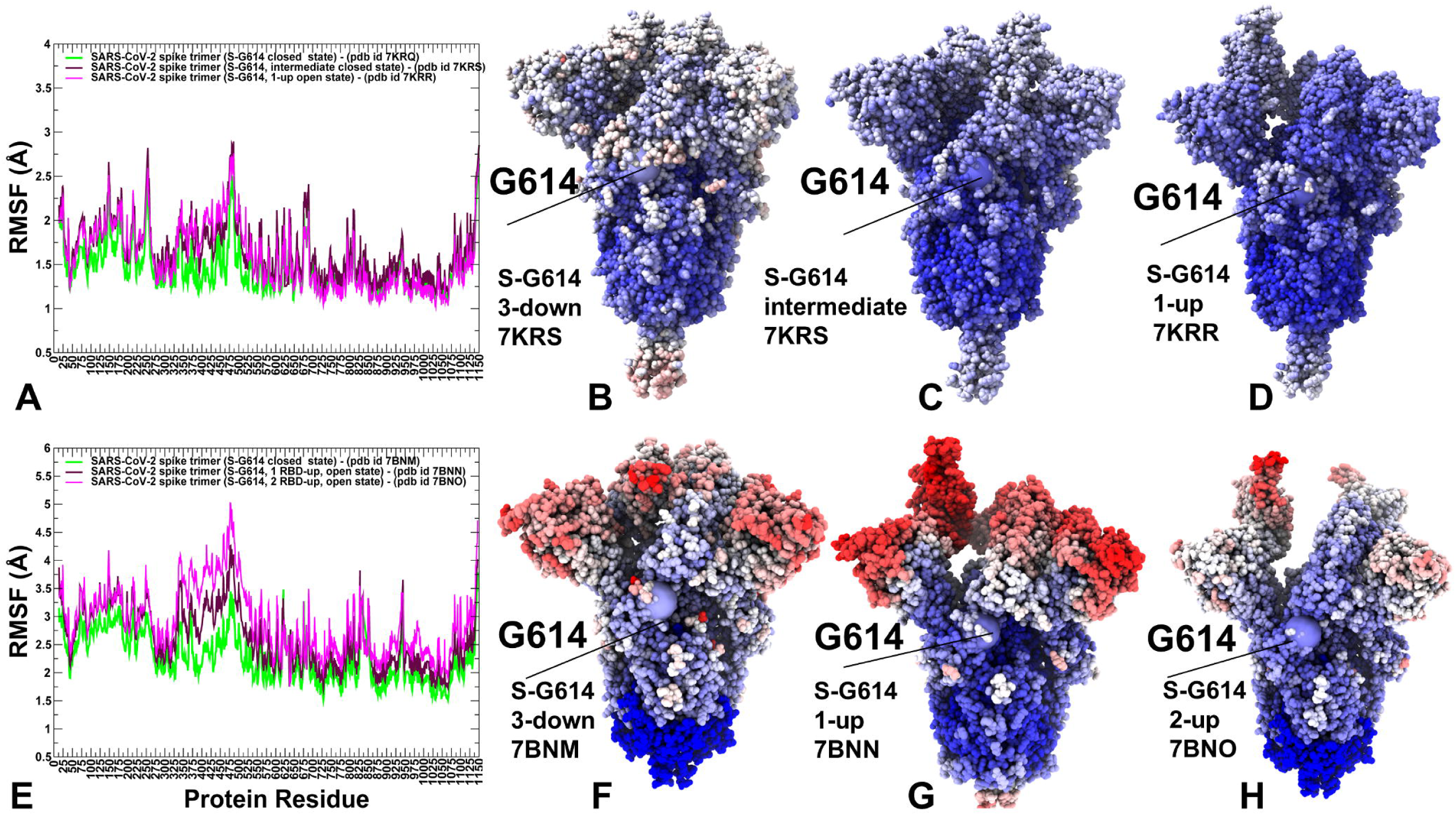
CG-CABS conformational dynamics profiles of the SARS-CoV-2 S-G614 structures. The S1 regions include NTD (residues 14-306), RBD (residues 331-528), CTD1 (residues 528-591), CTD2 (residues 592-686), UH (residues 736-781), HR1 (residues 910-985), CH (residues 986-1035), CD (residues 1035-1068), HR2 (residues 1069-1163). (A) The root mean square fluctuations (RMSF) profiles from simulations of the cryo-EM structures of SARS-CoV-2 S-G614 in the closed state, pdb id 7KRQ (green), in the intermediate state, pdb id 7KRS (red) and 1 RBD-up open form, pdb id 7KRR (blue). (B-D) Structural maps of the conformational profiles for the S-G614 closed state, the intermediate state and 1 RBD-up open form respectively. (E) The RMSF profiles of the cryo-EM structures of SARS-CoV-2 S-G614 in the closed state, pdb id 7BNM (green), 1 RBD-up state, pdb id 7BNN (red) and 2 RBD-up open form, pdb id 7BNO (blue). (F-H) Structural maps of the conformational mobility profiles for these S-G614 structures. The position of mutational site D614G is shown in spheres. The structures are in sphere-based representation rendered using UCSF ChimeraX package^146^ with the rigidity-to-flexibility sliding scale colored from blue to red.

We first performed simulations of the S-G614 mutant structures in the locked closed, intermediate and open forms (pdb id 7KRQ, 7KRS, and 7KRR respectively) solved by Chen and colleagues.^61^ These cryo-EM structures of the full-length S protein in the prefusion conformation discovered an ordered conformation of ∼25-residue FPPR segment that is typically disordered in other structures, demonstrating that RBDs can be locked in the tightly packed and rigid closed form (Figure 1A, D). The reported S-G614 mutant structures revealed structurally ordered 630 loop (residues 617 to 644) and FPPR region (residues 823 to 862). In the closed form of the S-G614 mutant, the FPPR segments (residues 823-862) and 630 loops are fully ordered (Figure 1A,D). In the intermediate conformation with a partly open single RBD, the 630 loops are disordered in the RBD-shifted module and FPPR remain fully ordered (Figure 1B, E).

Conformational dynamics analysis showed a considerable difference in the mobility profiles for these three conformational states of the S-G614 mutant (Figure 2A). Indeed, we found that the closed S-G614 trimer was generally stable, showing moderate but widely-spread fluctuations in the NTD and RBD regions (Figure 2B). Intriguingly, for the intermediate S-G614 mutant form, there is only marginal increase in the mobility of the S1 subunit, particularly RBD regions, while the S2 regions become more stable (Figure 2A,C). We observed that the intermediate form may be characterized by tight packing in the S2 regions and softening in the S1 subunit, which may be relevant for consolidation of the inter-protomer hinges typically formed at the borders of highly stable and more flexible regions (Figure 2C). Strikingly, a comparative analysis of structural maps of conformational mobility highlighted these changes, showing the increased stabilization of the partially open S-G614 trimer (Figure 2B-D). For the 1 RBD-up form of the S-G614 trimer (Figure 1C, F), we detected that only RBD and NTD regions remain fairly mobile (Figure 2A, D). These results are consistent with the notion that D614G mutation may induce the greater stability of the open conformations in which the remaining two closed RBDs are locked in their respective positions by the ordered 630 loops. Hence, the conformational dynamics analysis showed that the global mobility pattern is greatly influenced by structural constraints provided by the ordered 630 loop and FPPR segment. By highlighting the mutational D614G site, we noticed that this position may be located near the inter-domain hinges that form conformationally stable anchor regions that control global movements of the highly mobile RBD and NTD domains.

We also investigated conformational dynamics of another group of the closed (pdb id 7BNM), 1 RBD-up open (pdb id 7BNN) and 2 RBD-up (pdb id 7BNO) conformations of the S-G614 protein. The construct for the D614G mutant was based on the furin-uncleavable version of the SARS-CoV-2 spike protein ectodomain with a set of stabilizing mutations (R682S, R685S, K986P, and K987P).^63^ Notably, in theses S-G614 trimer conformations the 630 loop, the FPPR region (residues 828-853) and the furin cleavage site were disordered. It can be seen that all functional forms of the S-G614 mutant were fairly mobile, including the closed conformation (Figure 2E). In the closed state, the NTD regions, RBD (residues 331-528) and CTD1 (residues 528-591) corresponded to dynamic regions in the S1 subunit. While we observed the progressively increased fluctuations of the RBD residues in the 1 RBD-up and 2 RBD-up forms, the conformational mobility in the closed state was also evident (Figure 2E). The inter-domain regions and interfacial CTD1 residues linking S1 and S2 subunits showed an appreciable mobility level. Structural maps of conformational mobility displayed significant softening near the inter-protomer regions and at the S1/S2 interfaces (Figure 2F-H). We observed that G614 and the surrounding region displayed the increased conformational variability which may reflect structural weakening of the inter-protomer contacts and a wider topology of the S-G614 structures. These dynamic features may facilitate conformational transitions from the closed to a partially and fully open forms of S-G614 protein. The increased mobility of the S-G614 trimer in the fully open form could be seen from a comparison of structural mobility maps of the S-G614 structures (Figure 2F-H). It appeared that in the fully open form, S-614 trimer can experience larger fluctuations and partial expansions not only in the S1 subunit but also in the S2 regions that become more plastic and amenable for large functional changes. Hence, our results are consistent with structural studies^63^ showing that S-G614 mutant can assume a wide range of conformations, particularly increasing plasticity of the closed form, which may facilitate subsequent structural rearrangements to the open form.

We also simulated a group of the S-G614 open structures that represent three distinct 1 RBD-up substrates and two different 2 RBD-up stats characterized by different angle of the upward rotation with respect to the hinge region.^65^ These structures did not show significant conformational rearrangement around the interprotomer interaction site between G614 and T859 but revealed partially disordered K854 loop^65^ (Figure 3).

**Figure 3.**
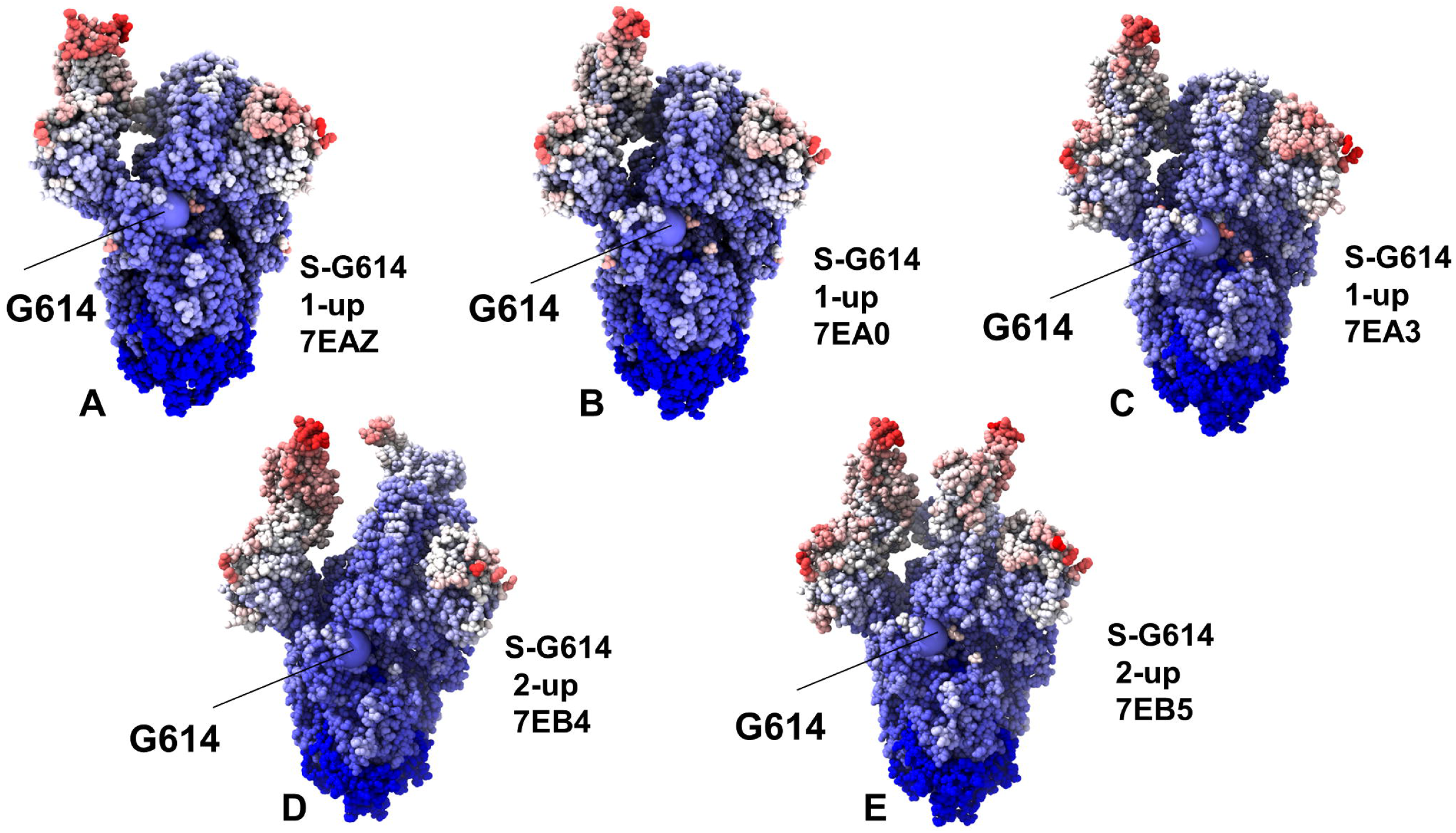
Structural maps of the conformational mobility profiles for the SARS-CoV-2 S-G614 1 RBD-up states (pdb id 7EAZ, 7EB0, 7EB3) (A-C) and 2 RBD-up states (pdb id 7EB4, 7lEB5) (D,E). (E) The position of mutational site D614G is highlighted in spheres and annotated. The structures are in sphere-based representation rendered using UCSF ChimeraX package^146^ with the rigidity-to-flexibility sliding scale colored from blue to red.

Structural maps of conformational mobility profiles obtained from CG-CABS simulations of these states revealed a fairly rigid S2 subdomain, mobile NTD and RBD regions for the 1-up protomer and the appreciable flexibility for one of the closed-down protomers (Figure 3A-C). This may reflect the increased propensity of 1 RBD-up conformations priming one of the closed protomers for the transition to the open state. The increased population of the RBD-up conformations and plasticity of the multiple open states may allow for specific modulation of the RBD motions and greater accessibility to the ACE2 host receptor.

The obtained conformational mobility profile for multiple 1 RBD-up substates is relatively similar, showing the rigidity of the S2 regions and mobility of the RBD-up regions (Figure 3A-C). It was also observed that the region harboring G614 site can bridge rigid and more flexible regions at the S1/S2 interface, with the site of mutation residing in stable local structural environment. These results suggest that modulation of local flexibility near the G614 position may affect the interfacial hinge regions and global movements of the RBD regions. Structural maps of the mobility profiles for 2 RBD-up states highlighted the increased plasticity of the S-G614 conformations in which both S1 and S2 subunits become “softer” and may experience significant fluctuations at the inter-protomer and S1/S2 interfacial regions (Figure 3D,E).

As a result, the hinge regions could migrate away from their positions, making the site of mutation and its local environment more flexible and decoupled from motions in the RBD regions. Hinges are the structurally rigid regions that link higher flexibility regions and may control their relative functional motions. Previous computational studies suggested that an enzyme evolves with subtle changes in dynamics and function, through a series of hinge-shift mutations by altering distinct, flexible positions with rigid positions. Our analysis indicated that changes in the inter-protomer hinge regions in the 2 RBD-up conformations and the increased conformational plasticity in the S1 subunit can further increase the diversity of receptor-accessible RBD conformations and promote association with the host receptor. Given that all considered structures may represent a spectrum of available conformational states for the S-G614 mutant, we infer that the S-G614 protein may sample a range of conformational states that promote frequent transitions to the open form.

### Functional Slow Modes of the SARS-CoV-2S-G614 Conformations Reveal Mediating Role of G614 in Communication of Hinge Regions Controlling Transitions between Open and Closed States

To identify hinge sites and characterize collective motions in the SARS-CoV-2 S-D614 and SARS-CoV-2 S-G614 structures, we performed PCA of atomistic reconstructed trajectories derived from CG simulations. Overall, the key functional signature of collective dynamics in these states are the preferences for NTD and RBD motions, suggesting that the conformational dynamics of these states can enable functional movements of RBDs to the erected, receptor-accessible conformation (Figure 4). Although functional slow modes follow a generally similar pattern for other group of S-G614 states, there are important differences reflecting the stability of the closed and open forms. The conserved hinge regions in the closed and open forms of the S-D614 protein that can regulate the inter-domain movements between RBD and NTD as well as the relative motions of S1 and S2 regions. We first compared the slow mode profile averaged over the first three lowest mode for the constrained S-G614 conformations (Figure S2). The major hinge positions are located at F318, F592, A570, I572, Q613, G614 and Y855 residues. These sites are specified in the distributions of slow modes (Figure S2) and emerged as immobilized islands in both the closed and open states. Notably, A570 and F592 hinge residues are situated near the inter-domain SD1-S2 interfaces and could act collectively as regulatory switch centers governing the population shifts between closed and open forms. Notably, for both 1 RBD-up and 2 RBD-up open conformations we observed significant functional displacements of at least two RBDs which suggests a significant degree of functional plasticity of the RBDs in the open conformations with the ordered 630 loop and FPPR region (Figure S2).

**Figure 4.**
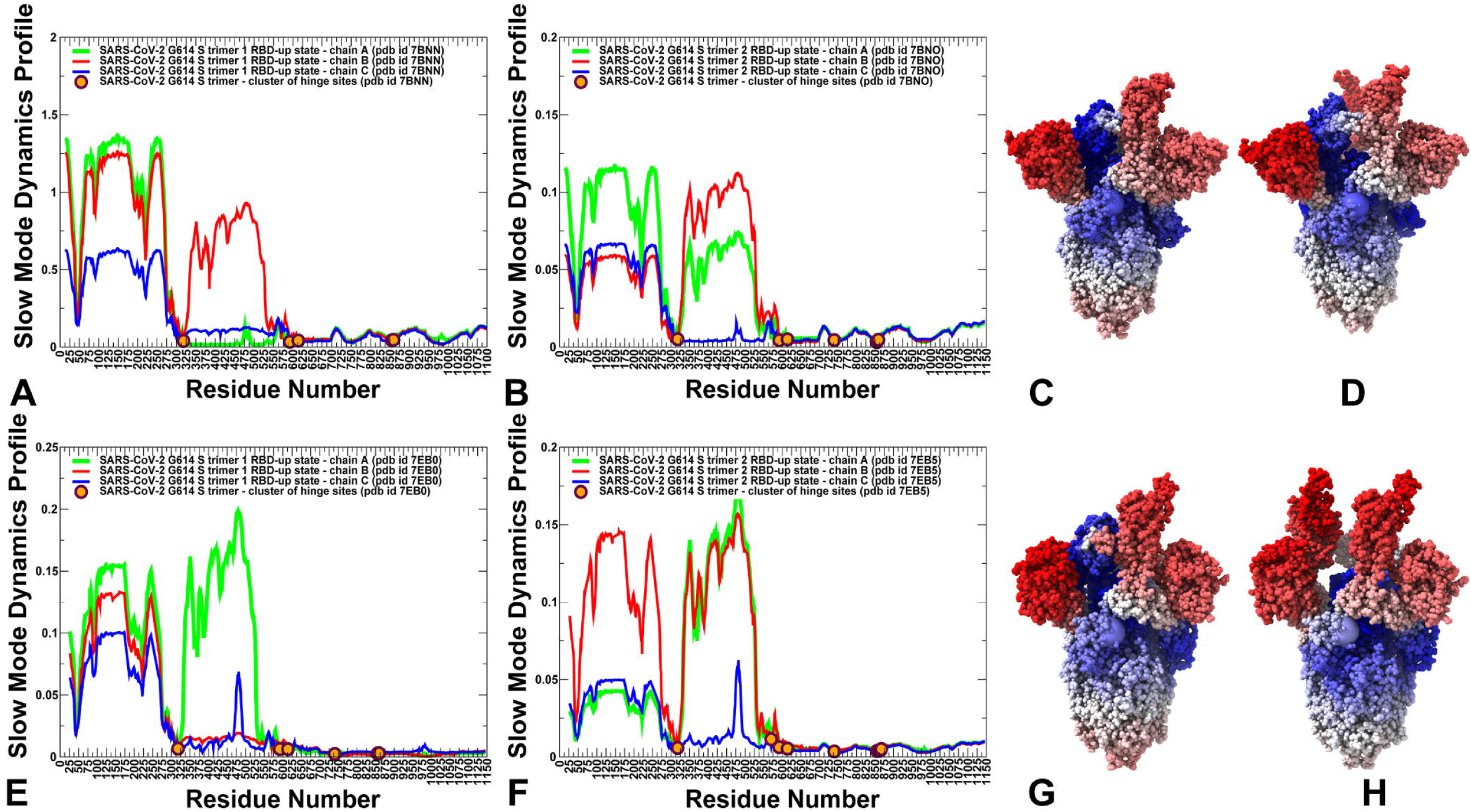
Functional dynamics of the SARS-CoV-2 S S-G614 trimer structures in the open form. The essential mobility profiles are averaged over the first three major low frequency modes. The essential mobility profiles for the SARS-CoV-2 S-G614 in the 1 RBD-up form, pdb id 7BNN (A), 2 RBD-up conformation, pdb id 7BNO (B). The profiles for protomer chains A, B and C are shown in green, red and blue lines respectively. The positions of the hinge sites forming the inter-protomer cluster F318, A570, F592, G614, M740, Y855, and T859 are shown along the profiles in filled orange-colored circles. Structural maps of the essential mobility profiles for the 1 RBD-up form, pdb id 7BNN (C) and 2 RBD-up conformation, pdb id 7BNO (D). The essential mobility profiles for the SARS-CoV-2 S-G614 in the 1 RBD-up form, pdb id 7EB0 (E), 2 RBD-up conformation, pdb id 7EB5 (F). Structural maps of the essential mobility profiles for the 1 RBD-up form, pdb id 7EB0 (G) and 2 RBD-up conformation, pdb id 7EB5 (H). The structures are shown in ribbons with the rigidity-to-flexibility scale colored from blue to red.

The observed pattern of collective motions in the S-G614 mutant would imply that the RBD regions may experience functional displacements in the intermediate form that encourage movements toward the open form. At the same time, once the S-G614 trimer attains the 1-up open form, the closed protomers become rigidified with their RBD movements being largely curtailed (Figure S2). As a result, unless open RBD becomes fully engaged by the ACE2 receptor, this may arguably prompt the open protomer to return to the down conformation as was proposed in structural studies.^61^

A generally similar pattern was seen in the slow mode analysis of the multiple S–G614 open conformations (Figure 4). The analysis of slow modes for the flexible open conformations^63^ (Figure 4A,B) showed significant displacements of NTD and RBD regions. In particular, we noticed that RBD motions in the 2 RBD-up state become more pronounced. The slow mode profiles and structural maps of functional movements revealed that G614 is situated near major hinge clusters separating stable and flexible regions and can be involved in orchestrating RBD movements (Figure 4C,D).

Indeed, the mutational site is located in the immobilized structural region of the SD2 domain that together with β strand formed by residues 311–319 may correspond to a hinge center governing motions of NTD and RBD, as well motions in S1 from the S2 subunit. The flexible 28-residue N2R linker (residues 306–334) connects the NTD (residues 27–305) and RBD (residues 335–521) and stacks against the SD1 (residues 529–591) and SD2 (residues 592–697) The residues 311–319 of the linker that associate with the SD2 subdomain remained immobilized in slow modes forms a hinge region separating the mobile NTD and RBD, as well as isolating the motions in S1 from the S2 subunit. The broad hinge regions are formed by residue clusters K310-F318 that mediates NTD-RBD motions and F592-V618 that control RBD-S2 motions (Figure 4).

The slow mode profiles obtained multiple 1 RBD-up and 2 RBD-up substates of S-G614 structures^65^ have a similar shape, featuring large displacements of NTD and RBD regions for the ACE2-accessible protomers (Figure 4 E,F). Of notice are appreciable displacements for the RBD-down domain in the 2 RBD-up structure (Figure 4F), particularly pointing to a peak aligned with E484 residue in this protomer. This indicated that the highly flexible 2 RBD-up states could promote the increased mobility of the closed RBD protomer and prime S-G614 for subsequent transition to 3 RBD-up state that maximizes the exposure of the S protein to binding with ACE2. Structural maps for these states highlighted the broadened area of functional motions beyond highly mobile NTD and RBD regions (Figure 4 G,H). In addition, we noticed a somewhat weakened level of immobilization for the G614 position, indicating partial reorganization of the hinge clusters in these states. These results suggested that the D614G mutation confers the increased structural flexibility in the closed state which may promote the exchange between the open and closed forms and the increased exposure of the RBDs for interactions with ACE2. In addition to major hinge points centered on F318 and A570/F592, a more detailed analysis of the slow mode profiles also revealed presence of local minima corresponding to residues D737/M740 and K854/F855 (Figure 4). This is evident for the functional motion profiles of the constrained S-G614 states and 2 RBD-up conformations (Figure 4B,F). Hence, conformational changes in the S protein may be orchestrated through a cross-talk of several hinge centers that could couple NTD-RBD and RBD-S2 motions.

Intriguingly, structural analysis of the ensemble-averaged S-G614 states revealed an important role of G614 site in mediating communication between these hinge clusters (Figure 5). The cluster of hinge sites formed by K854, F855, F592, P589, D737 and M740 is fairly well-packed and remains intact in both closed and open S-G614 states (Figure 5). In the ensemble of structurally constrained S-G614 conformations with the ordered 630 loop, G614 is weakly linked with the hinge cluster through the key hinge position F592 (Figure 5A,B). The presence of structurally ordered loop may limit lateral movements near G614 position and provide only suboptimal interactions with F592. Nonetheless, the strategic position of G614 could allow for communication between the hinge cluster (K854, F855, F592, P589, D737 and M740) and other hinge center (F592-V618) that control S1-S2 motions. In more flexible open S-G614 conformations with disordered 630 loop, G614 can experience more significant lateral displacements by approaching and departing from F592-centered hinge cluster (Figure 5C,D). As a result, we argue that G614 site may serve as a versatile regulatory switch point that links or decouples the network of hinge clusters. In this mechanism, the disorder-order transitions of the flexible 630 loop could allosterically modulate G614 couplings with the hinge clusters and therefore contribute to the control over collective motions of the S1 subunit.

**Figure 5.**
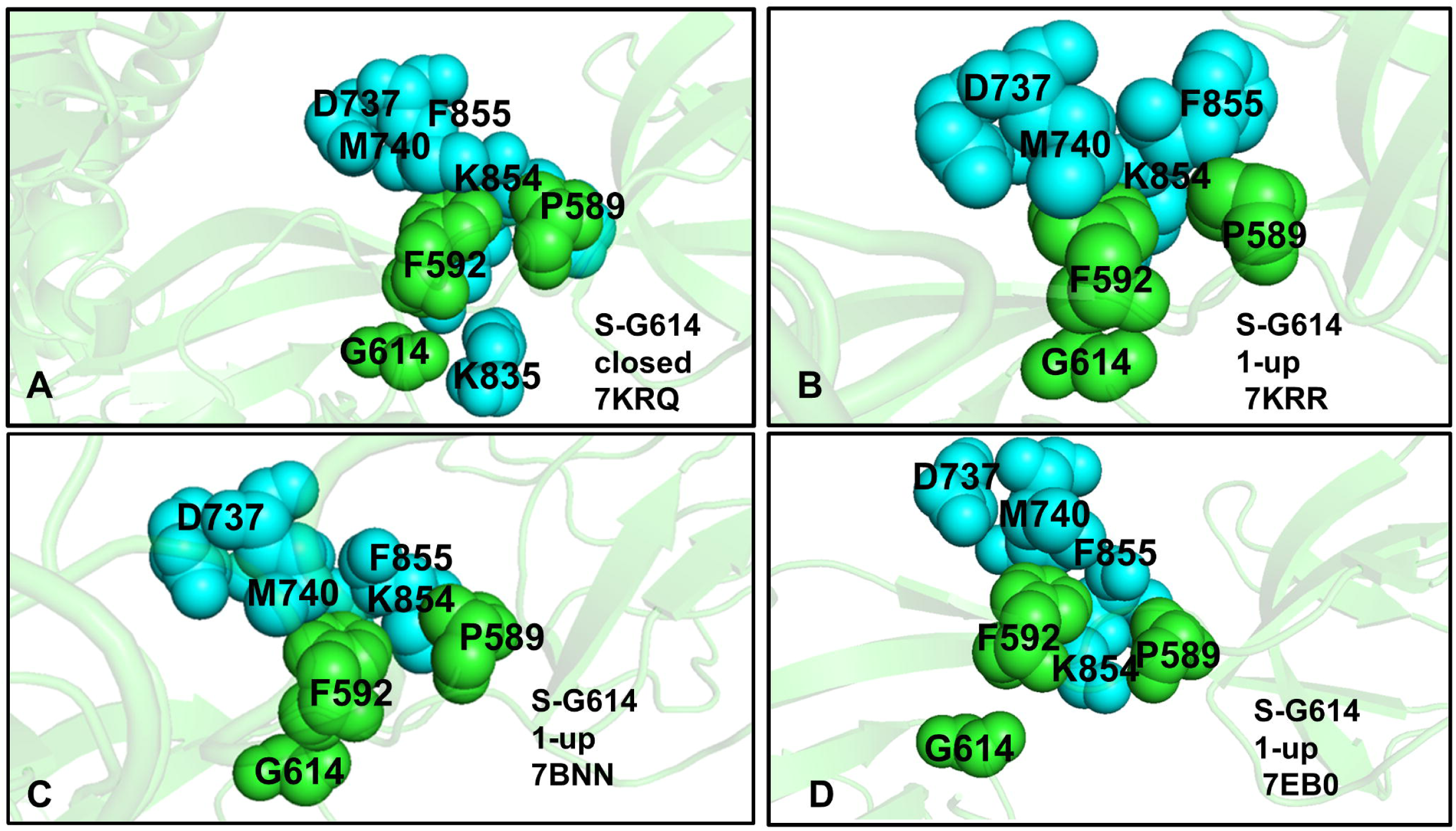
Structural analysis of the ensemble-averaged S-G614 states in the closed and open forms. (A) A close-up of the interaction cluster between G614 and hinge region formed by F592 and P589 of the same protomer (in green) and K835, K854, F855, M740, D737 and F855 of another protomer (in cyan) in the closed form (pdb id 7KRQ). (B) A close-up of the interaction cluster between G614 and hinge region in the 1 RBD-up open form (pdb id 7KRR). (C) A close-up of the interaction cluster between G614 and hinge region in the 1 RBD-up open form (pdb id 7BNN). (D) A close-up of the interaction cluster between G614 and hinge region in the 1 RBD-up open form (pdb id 7EB0). Note that G614 is linked to the hinge cluster through a critical hinge hub F592.

### Local Frustration Patterns in the Ensembles of SARS-CoV-2 S-G614 States: Mutational Frustration Neutrality of Variant Sites and Modulation of Frustration in the Open States

We employed the conformational ensembles of the multiple conformational states of the S-G614 trimer to estimate the average local frustration profiles of the S residues and quantify the relationship between structural plasticity and degree of mutational frustration. This analysis is based on scanning conformational ensembles by local frustratometer^147, 148^ which computes the local frustration index based on the contribution of a residue or residue pairs to the energy in a given conformation as compared to what it would contribute in decoy conformations. In this approach by performing mutations and changing local environments, the mean and variance of the energies of possible decoys relative to the native conformation are evaluated.^147–151^ Previous studies by Wolynes and colleagues discovered that minimally frustrated residues are typically enriched in the rigid protein core, while a large fraction of residues are characterized by neutral frustration, i.e. these positions residue near the center of the distribution of possible energies in decoy states.^149–151^ At the same time, it was found that residues near the binding sites the proteins are enriched in highly frustrated interactions.

We first examined the distribution of local mutational frustration in different substates of the S-G614 protein, particularly focusing on G614 position as well as sites targeted by common circulating variants in the RBD (K417, E484, N501) (Figure 6). The role of frustration in the RBD regions is particularly interesting in light of recent evidence that the disordered/highly flexible regions are important for mediating allostery and binding to multiple protein partners, where energetic frustration at the molecular interfaces may serve as a mechanism for allosteric regulation [96-99].

**Figure 6.**
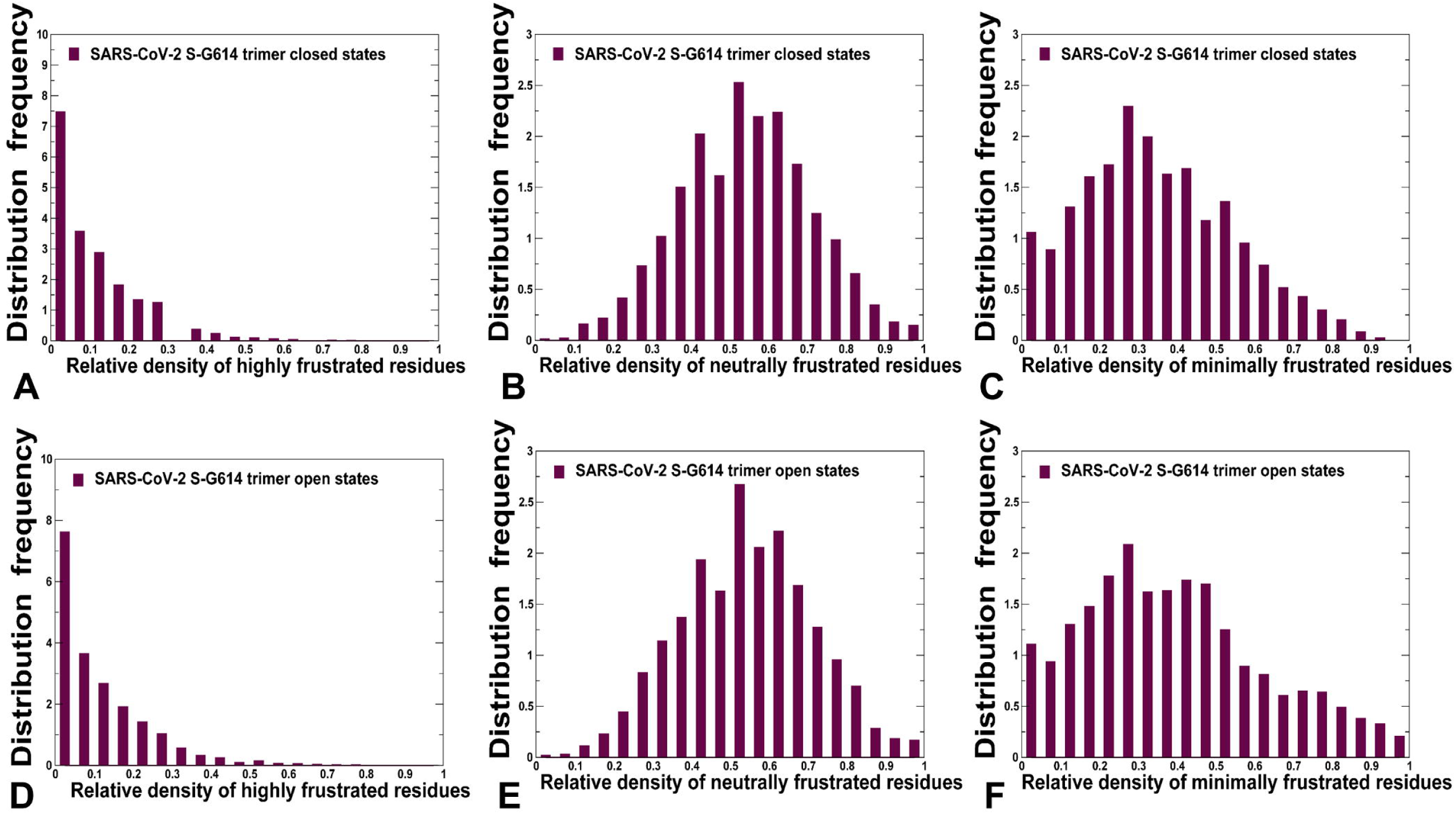
The distribution of local mutational frustration in the closed and open forms of the SARS-CoV-2 S-G614 variant. (A-C) The relative density of highly frustrated, neutrally frustrated and minimally frustrated residues respectively in the closed S-G614 conformations. The distributions were constructed by averaging computations over snapshots from simulations of the closed states (pdb ids 7KRQ and 7BNM). (D-F) The relative density of highly frustrated, neutrally frustrated and minimally frustrated residues respectively in the open S-G614 conformations. The distributions were constructed by averaging computations over snapshots from simulations of all 1 RBD-up and 2 RBD-up states (pdb ids 7KRR, 7BNN, 7BNO, 7EAZ, 7REB0, 7EB3, 7EB4, 7EB5).

First, we evaluated the cumulative distribution of local frustration index in the closed and open S-G614 states by averaging the residue-based local mutational frustration over dynamic ensembles of conformations (Figure 6). Interestingly, similar distributions of highly frustrated, neutral and minimally frustrated states respectively were seen in the closed (Figure 6A-C) and open forms (Figure 6D-F). Notably, the population of highly frustrated conformations is relatively minor, but we observed a longer density tail of highly frustrated values for the distribution in the open states (Figure 6D). This may reflect a considerable conformational heterogeneity of the multiple open conformations where the frustrated sites correspond to highly exposed positions in the NTD and RBD. Importantly, we observed the dominant population of neutrally frustrated sites in both closed and open forms as well as a significant density of minimally frustrated regions. A relatively small fraction of highly frustrated positions as compared to neutral and minimally frustrated residues is an important observation of this analysis. This suggested that while the exposed RBD and NTD regions in the open states are fairly flexible, the degree of mutational frustration and adaptability in these positions may be moderate to enable structural stability and binding with the host receptor.

A comparison of local mutational frustration indexes for key functional positions including sites of variants (K417, E484, and N501), hinge position A570 and G614 revealed interesting shifts between the closed and open forms of S-G614 trimer (Figure 7). The high frustration index for these positions is uniformly small and similar in both closed and open conformations (Figure 7A,D). While this may be expected for hinge positions A570 and G614, it was somewhat revealing for the RBD positions, especially mobile E484 and N501 residues. It is of some notice that the relative high frustration density for G614 site becomes even smaller in the open form, indicating stabilizing effect of G614 residue in the receptor-accessible states (Figure 7A,D). The relative density of neutral frustration is pronounced for all sites, especially for N501 and G614 positions, showing no appreciable differences between the closed and open states (Figure 7B,E).

**Figure 7.**
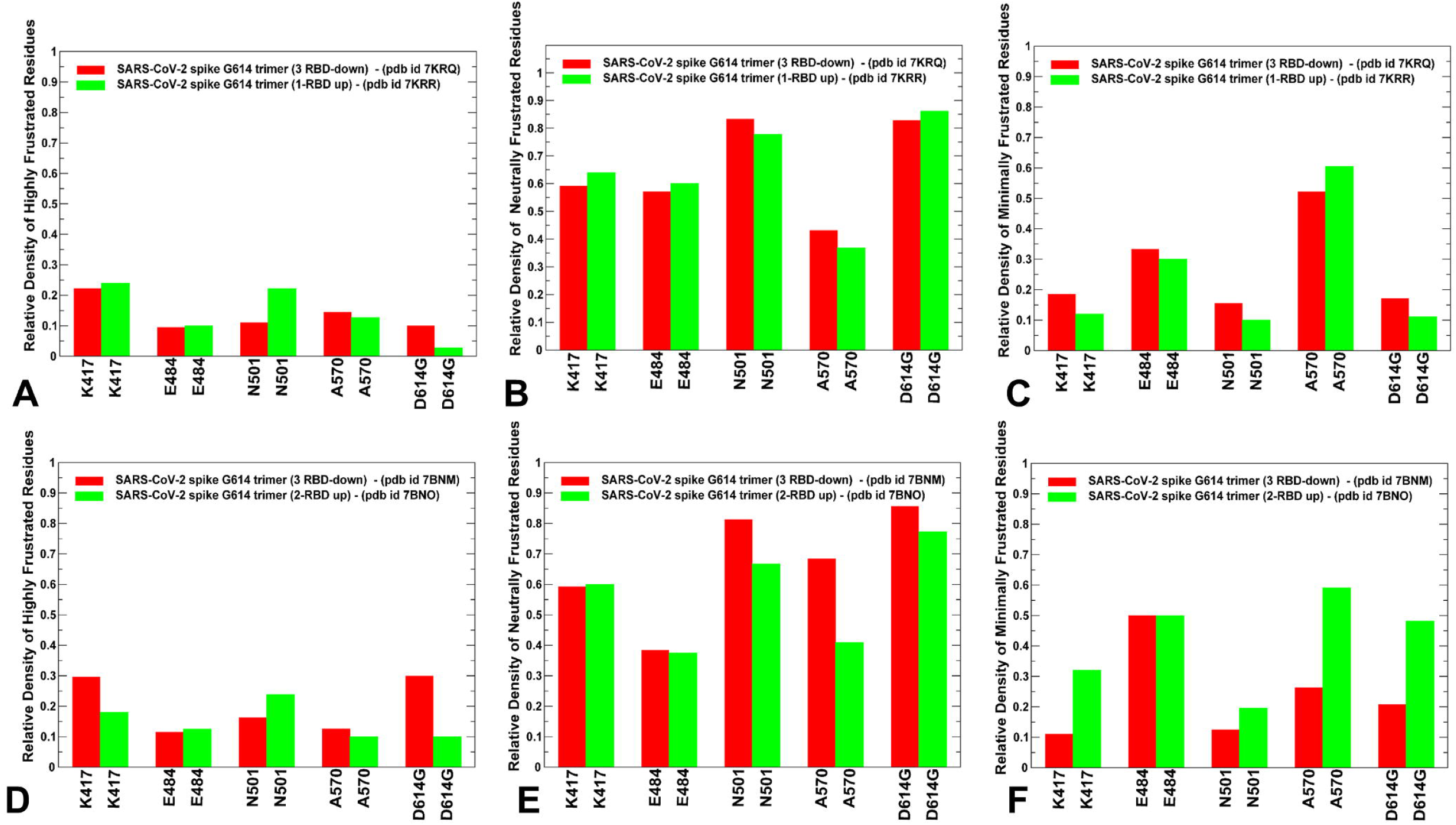
A comparison of the ensemble-averaged local mutational frustration between closed and open forms for sites of mutational variants and hinge positions (K417, E484, N501, A570, and G614) (A-C) The relative density of highly frustrated, neutrally frustrated and minimally frustrated residues respectively for the functional sites in the closed state (pdb id 7KRQ) and 1 RBD-up open state (pdb id 7KRR). (D-F) The relative density of highly frustrated, neutrally frustrated and minimally frustrated residues respectively for the functional sites in the closed state (pdb id 7BNM) and 2 RBD-up open state (pdb id 7BNO).

This suggested that sites of important circulating variants located in the flexible RBD regions showed a preponderance towards neutral mutational frustration. An appreciable density of minimal frustration for E484, A570 and especially G614 positions was also found (Figure 7C,F). In particular, we observed the increased minimal frustration of G614 in the ensemble of open conformations. Hence, the interactions of this key functional residue may become less frustrated in the open form as compared to the closed states. This is an interesting observation, particularly given that the minimally frustrated interactions are often excluded from the vicinity of binding residues in globular proteins [94,95]. According to our results, the conformational plasticity of the RBD-up ensembles can allow for thermal rearrangements of the inter-protomer contact and repacking near G614 position, providing means for neutral-to-minimum frustration level in the mutational site. We argue that this allows for moderately suboptimal inter-protomer interactions leading to multiple open substates, each displaying specific frustration patterns, which can be exploited to control binding with the ACE2 and to multiple partners.

According to our data, the relative density of neutral frustration and minimal frustration can be similar for G614 position in the ensemble of open states. By modulating the frustration level and local dynamic environment around the A570/G614 hinge region, D614G mutation may modulate the thermodynamic equilibrium between the closed and open states as well as level of functional plasticity in the open substates required for allosteric function and/or binding.

### Mutational Scanning of Protein Stability the SARS-CoV-2 S-614 Conformational States Reveals Energetic Effects of the D614G Mutation

We employed the equilibrium ensembles generated from CG-CABS simulations of the SARS-CoV-2 S protein structures to perform mutational scanning analysis of the S-G614 proteins (Figure 8). The questions addressed in this analysis are to examine the effect of the D614G mutation on different functional forms of the S-G614 protein and quantify the mutation-induced mechanism of S protein stabilization. The protein stability ΔΔG changes were computed by averaging the results of computations over 1,000 samples obtained from simulation trajectories. We constructed mutational sensitivity heatmaps for the major sites of circulating variants in the RBD (K417, E484, and N501) and G614 position in the S-G614 closed and open trimer forms. Mutational cartography revealed several interesting trends and highlighted the energetic preferences of G614 site (Figure 8). Interestingly, for the constrained S -G614 states, modifications in this position generally resulted in appreciable destabilization changes, while protein stability changes are only moderately perturbed by mutations in K417, E484 and N501 sites. Despite different level of structural plasticity in the examined closed and open states, mutational scanning map showed that the local structural environment around G614 is still fairly adaptable (Figure 8). Indeed, while modifications in the G614 position generally produced destabilizing free energy changes, the magnitude of these changes was relatively moderate. Notably, the magnitude of mutation-induced changes was smaller in the flexible open S-G614 states as compared to more constrained conformations (Figure 8). These findings are consistent with the mutational frustration analysis, indicating that the local environment near G614 is neutrally-to-minimally frustrated in the open conformations. In particular, the structural map showed a significant degree of energetic tolerance to G614 mutations in one of the 1 RBD-up open states (pdb id 7EB0) and 2 RBD-up conformations (Figure 8). At the same time, mutational scanning maps for the more dynamic S-G614 states displayed a pattern of moderate destabilization or neutral energetic changes for the RBD positions K417, E484 an N501 (Figure 8). The mutational cartography analysis revealed that a moderate level of energetic frustration and suboptimal interactions are characteristic not only of the sites of circulating variants located in the RBD but also for the D614G position. These energetic results are consistent and reciprocate the mutational frustration analysis, showing that the disordered and structured regions are enriched in neutrally (moderately) frustrated interactions. Our analysis suggested that D614G mutation may elicit sub-optimal interaction contacts and moderate level of frustration near the inter-protomer interfaces and hinge clusters.

**Figure 8.**
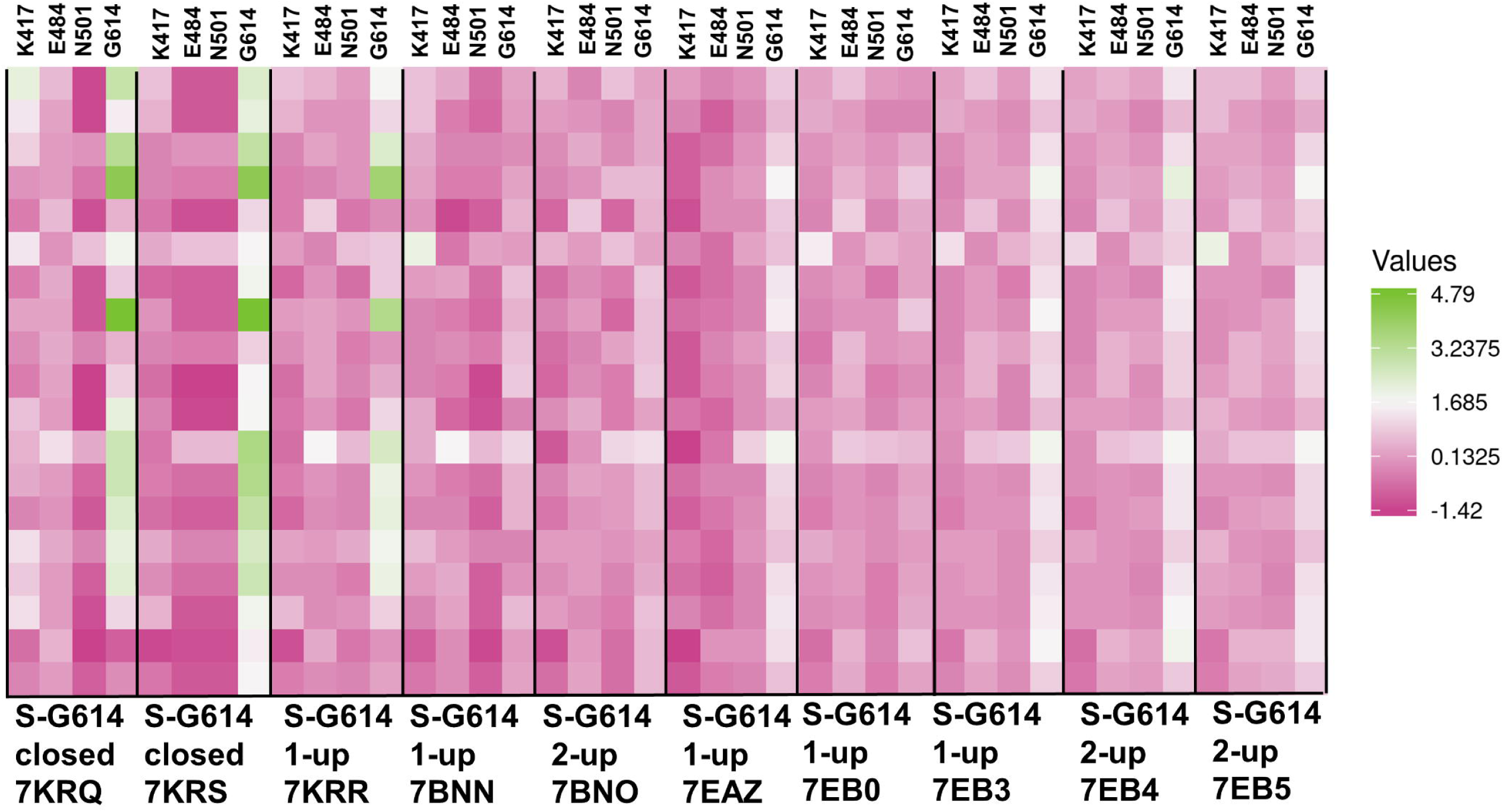
Ensemble-based mutational profiling of the SARS-CoV-2 S-G614 stability. The mutational scanning heatmaps for the multiple SARS-CoV-2 S-G614 states at positions K417, E484, N501, and G614.The heatmaps show the computed binding free energy changes for 19 single mutations on the sites of variants. The squares on the heatmap are colored using a 3-colored scale – from pink to white and to green, with green indicating the largest unfavorable effect on binding.. The standard errors of the mean for binding free energy changes were based on 5 independent CG trajectories for each of the S-G614 states and different number of selected samples from a given trajectory (500, 1,000 and 2,000 samples) are ∼ 0.15-0.22 kcal/mol using averages over different trajectories and ≤ 0.15 kcal/mol from computations based on different number of samples from a single trajectory.

We argue that the energetic frustration near G614 could allow for communication between hinge clusters and allosterically modulate the functional transitions of the S1 subunit. This landscape-based mechanism of mutation-induced energetic frustration in the S protein may result in the greater adaptability and the emergence of multiple conformational substates in the open form. The results also suggested an interesting interplay between mutation-induced protein stability and local frustration patterns. It is particularly instructive to interpret these relationships in the context of allosteric regulation models centered on the energetic frustration that could emerge at the inter-domain interfaces.^152–155^ In our case, the D614G mutation may introduce the level of energetic frustration and conformational plasticity near the inter-protomer and inter-domain interfaces that would allow for efficient modulation of allosteric couplings between the hinge clusters without compromising the structural integrity of the S protein. Through this frustration-inspired allosteric mechanism, the D614G mutation can impose allosteric control over functional movements and conformational diversity of the RBD regions.

A more detailed analysis of mutational scanning in the constrained S-G614 states showed significant destabilization upon modifications in both closed and open states. Interestingly, however, hydrophobic modifications to F/W may produce marginally improved stability (Figure S3). The stabilization predicted by these mutations may be due to the hydrophobic effect and improved non-specific packing near the mutational site. Interestingly, the magnitude of destabilization changes was larger in the intermediate close state, particularly for substitutions that change the size and chemical nature of the amino acid such as G614D/E/K (Figure S3). Somewhat smaller but still significant destabilization changes upon G614 mutations are also detected in the 1 RBD-up open state. These results suggested that conformational plasticity is present even in the constrained S-G614 states which is consistent with the increased flexibility of the S protein and preferences for the open state induced by G614. These results also corroborated with the frustration analysis, suggesting a moderate degree of frustration in the G614 position for open states. The range of destabilization free energy changes induced by mutations in the G614 position was considerably reduced in more flexible open structures determined by Benton and colleagues^63^ (Figure S4). While the destabilization changes remain relatively pronounced in the closed form, the protein stability energy differences were very small in the 1 RBD-up and 2 RBD-up open states (Figure S4). The range of free energy ΔΔG changes where some modifications could even improve stability (G614D and G614N) was less than 1 kcal/mol, suggesting that these wide and mobile open conformations feature a considerable degree of conformational plasticity and local frustration near the mutational site. Accordingly, mutational tolerance of the S-G614 structures could facilitate transitions to the ensemble of open conformations with significant degree of conformational heterogeneity. We also detailed protein stability changes induced by G614 mutations in the ensemble of multiple conformational substates of the S-D614^65^ (Figure S5). For these conformations, we generally observed moderate destabilization changes ΔΔG ∼ 1.0-1.5 kcal/mol. In particular, we noticed that modifications G614E/D/K and G614 substitutions to larger hydrophobic residues were highly unfavorable (Figure S5). Hence, these S-G614 open substates featured only moderate degree of local frustration and signs of minimal frustration as the local interactions near mutational site were more optimal (Figure S5). These results supported our hypothesis that conformational plasticity of the RBD-up structures can allow for neutral-to-minimum frustration level near the mutational G614 site. The suboptimal inter-protomer interactions in the multiple open substates featuring moderate frustration can allow for modulation of functional movements and may promote conformational transitions to the ensemble of open S-G614 states.

### Dynamic Network Modeling and Ensemble-Based Community Analysis Identify Allosteric Interaction Networks in the SARS-CoV-2 Spike Mutants

Network-centric models of protein structure and dynamics used in the current study can also provide a complementary perspective to physics-based landscape models and allow for quantitative analysis of allosteric changes, identification of regulatory control centers and mapping of allosteric communication pathways. Using this framework, the residue interaction networks in the SARS-CoV-2 spike trimer structures were built using a graph-based representation of protein structures^124, 125^ in which residue nodes are interconnected through both dynamic^126^ and coevolutionary correlations.^127^ By employing perturbation-based network modeling, we computed ensemble-averaged distributions of the betweenness centrality based on average shortest path length (ASPL) change upon removing/mutating individual nodes (Figure 9). Using these parameters, we identify mediating centers of allosteric interactions in the SARS-CoV-2 mutant structures. The residue interaction networks were also divided into local interaction communities in which residues are densely interconnected whereas residues from different communities may be weakly connected through the inter-modular links.

**Figure 9.**
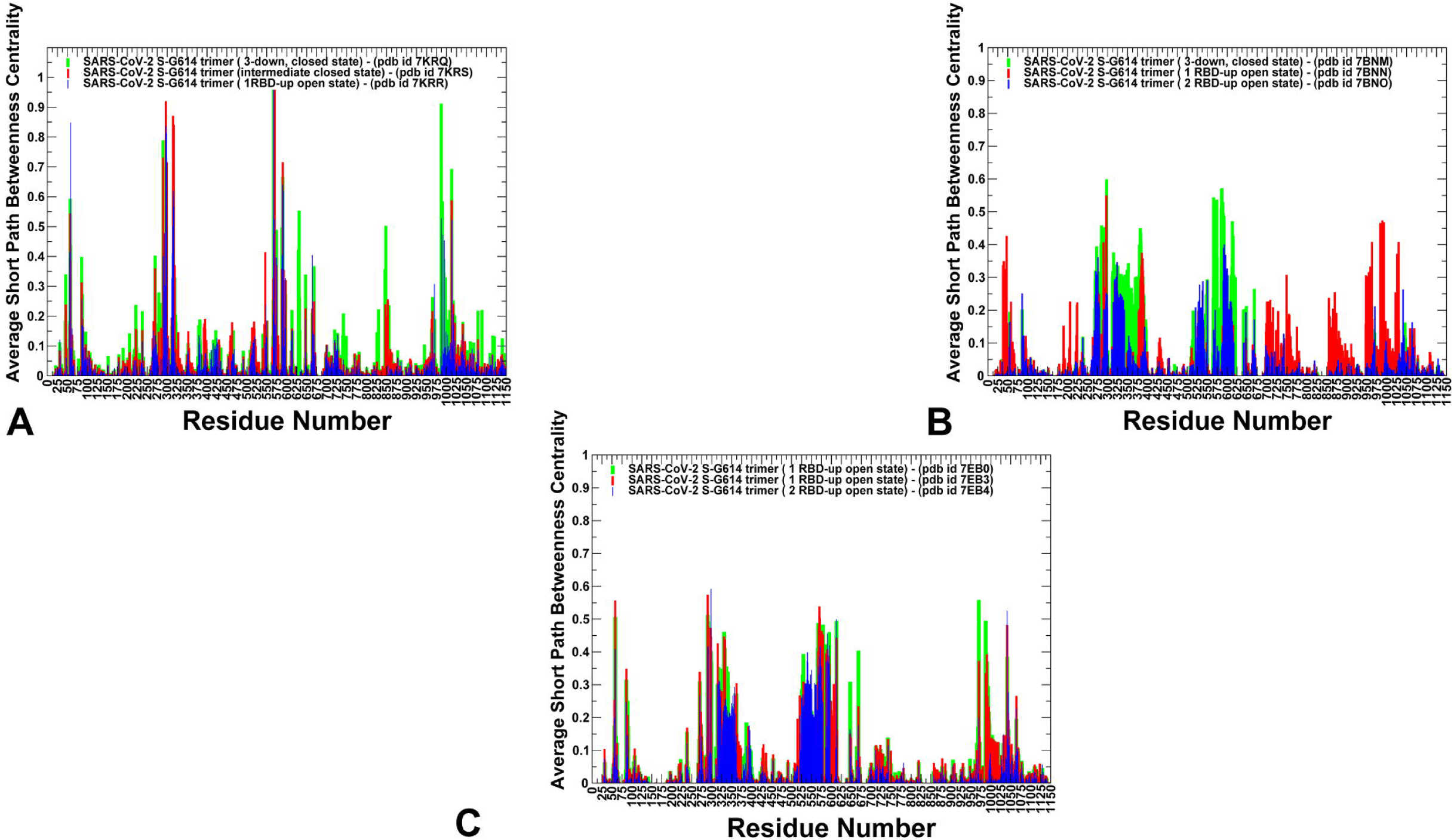
Network centrality analysis of the SARS-CoV-2 S-G614 trimers. (A) The ensemble-averaged short path betweenness centrality for the structure of SARS-CoV-2 S-G614 in the closed state, pdb id 7KRQ (green lines), the intermediate form, pdb id 7KRS (red lines) and open form, pdb id 7KRR (blue lines). (B) The ensemble-averaged short path betweenness centrality for the structure of SARS-CoV-2 S-G614 in the closed state, pdb id 7BNM (green lines), the 1 RBD-up open form, pdb id 7BNN (red lines) and 2 RBD-up open form, pdb id 7BNO (blue lines). (B) The ensemble-averaged short path betweenness centrality for the structure of SARS-CoV-2 S-G614 in the 1 RBD-up open substate form, pdb id 7EB0 (green lines), 1 RBD-up substate form, pdb id 7EB3 (red lines), and 2 RBD-up open form, pdb id 7EB4 (blue lines). The SARS-CoV-2 S mutant trimers are shown in ribbons. The residue-based profiles are shown for protomers of the S trimer in green, red, and blue lines. The betweenness centrality profiles based on ASPL change upon removing individual nodes are shown for the structures.

We first constructed the distribution of the short path betweenness profile for the structurally constrained S-G614 conformations (including closed, intermediate and partly open states) (Figure 9A). These S-G614 mutant structures that were solved with structurally ordered 630 loop (residues 617 to 644) and FPPR region (residues 823 to 862).^61^ We noticed that the network centrality profile displayed appreciable peaks in these regions (Figure 9A). These observations are consistent with structural studies indicating that the structured 630 loop in the S-G614 trimer can promote the tighter packing among three protomers and also stabilizes the couplings with CTD2 which inhibits the release of the N-terminal segment of S2 and blocking S1 dissociation.^61^ The short betweenness profiles for the more rigid S-G614 states showed sharp and narrow peaks in the CTD1 region corresponding to residues D568 A570, T572, and F592 in the CTD1 region (Figure 9A). Several other peaks along the network betweenness profile corresponded to hinge centers F318, V539, L570, I572, T547, F592, S750, I856 and L962 residues (Figure 9A). Together, high centrality clusters could mediate allosteric couplings between the 630 loop, CTD1 residues and RBD regions. Notably, these regulatory points are located near the inter-protomer and inter-domain contact interfaces. The emergence of sharp single peaks in the distribution aligned with mediating centers in the S-G614 trimer signaled those allosteric communications in the structurally constrained S-G614 closed and open states may proceed through a fairly narrow pathway connecting key mediating sites located in CTD1 regions. In network terms, this would allow for efficient and direct signaling but could imply a degree of vulnerability to mutational perturbations of key mediating centers.

A noticeable change in the density of mediating residues was seen for the centrality distributions obtained for more flexible S-G614 open states (Figure 9B). For these structures, we observed the broadening of the density peaks in the CTD1 region and in the flexible 28-residue N2R linker spanning residues 306–334 that connects the NTD (residues 27–305) and RBD (residues 335–521) and stacks against the SD1 (residues 529–591) and SD2 (residues 592–697) regions. Significantly, the bifurcated distribution featured fairly broad peaks for the N2R and CTD1 regions, suggesting that the conformational flexibility of the S-G614 open states would translate to a much broader and arguably more robust ensemble of communication paths in the S-G614 structure. Notably, for all S-G614 open conformations, the G614 position was associated with a pronounced distribution peak. A similar pattern was obtained from the network analysis of the multiple substates of the open S-G614 conformations (Figure 9C).

In these structures, we also observed two very broad and distinct peaks of the short path centrality aligned with N2R and CTD1 regions, strongly arguing for a broad ensemble of signal transmission routes in the conformationally mobile open S-G614 states. The observed network signatures imply that structurally rigid hinge positions in CTD1 (A570, T572, F592 and G614) are coupled with more flexible residues to form a broad “tube” of allosteric communication routes. This would provide a significant resiliency and robustness to signaling in the S-G614 structures, making the allosteric function of the S protein robust against perturbations and mutations. The results also suggested an important role to the N2R linker that connects the NTD regions to the RBD residues within a given protomer. Rather than just being a connector, this 28-residue linker could also function as a modulator of allosteric conformational changes that are critical for the ACE2 receptor engagement. The important conclusion of this analysis is that by inducing conformational heterogeneity of the open S form, the D614G variant allows for robust allosteric communications between the site of mutation, CTD1, NTD and RBD regions. In this scenario, allosteric signal initiated near the mutational site in the open state, would propagate via a broad “tube” of communication routes to the RBD regions, making allosteric interactions robust to perturbations and variations in the ensemble of the S conformations.

We also performed community analysis to monitor dynamic changes in the modular organization of the inter-protomer interfaces. The community-based analysis and modularity assessment of allosteric interaction networks is based on the notion that groups of residues that form local interacting communities are expected to be highly correlated and can switch their conformational states cooperatively. As a result, long-range allosteric interaction signals can be transmitted through a hierarchical chain of local communities on the dynamic interaction networks.

The community analysis of the S-G614 structures is used as proxy for measuring protein stability and communication efficiency in performing allosteric functions. This analysis provides also a global metric to measure changes in connectivity and interaction between subdomains and inter-protomer association in the SARS-CoV-2 S structures. In the closed S-G614 rigid trimer, the interfacial communities in the hinge region include A614-C855-C589-C592; A568-A574-C854; A614-A740-A857-B592; B775-B864-A665; A592-C737-C855 (Figure 10A,B). The hinge positions in the closed state anchor the inter-protomer communities A568-A-574-C854; B855-C589-C592; A614-A592-A589-B835-B854; A614-B740-B737-A592; and A592-C737-C855 (Figure 10A,B) and control propagation of allosteric signal between S1 and S2 subdomains through a fairly narrow channel formed by these communities in the CTD1 region. In the S -G614 trimer, G614 forms community with F592 and P589 of the same protomer, K835 and K854 of the adjacent protomer (Figure 10A,B). This rewiring allows for strong local coupling of G614 and hinge center F592 and provides a “stapling” lock for the rigid close conformation. This analysis provided an additional support to the notion that D614G can function as an important allosteric hub for mediating the increased flexibility in the SD1-S2 hinge regions. An emerging concept central to computational models of allostery is identification of conformational switch centers that mediate long-range communications and allosteric pathways.^156–158^ Our analysis suggested that these inter-protomer communities near hinge centers could function as such allosteric security switches that facilitate large scale transitions to the open form. We observed several common changes in the inter-protomer communities of the more flexible 1 RBD-up conformations (Figure 10C,D). First, G614 and P589 weaken their interactions with the communities A614-B855-C589-C592 and G614 becomes further decoupled from the community A614-B740-B737-A592 (Figure 10C,D). Second, we found that the local intra-protomer community formed by residues A570, D568 and D574 is tightly coupled to the network of the interfacial communities in the closed S-G614 trimer (Figure 10A,B), but becomes separated from the inter-protomer communities in the 1 RBD-up open form due to greater conformational variability in this region (Figure 10C,D). In the ensemble-averaged analysis, the weakened contacts between these communities become further exacerbated which leads to fragmentation of the major hinge cluster formed by F318, F592, A570, I572, G614 and Y855 residues. As a result, in the open S-G614 states, G614 site is community-coupled to F855 and M740 positons of the same protomer and hinge site F592 of the adjacent protomer. In addition, an important hinge center A570 becomes disengaged from the inter-protomer hinge cluster that may release some constraints on movements in the S1 regions and promote greater conformational flexibility of the open protomers. These observations from the community analysis are consistent with the local frustration profiling showing modulation of the frustration level and local dynamic environment around the A570/G614 hinge region, which may increase functional plasticity in the open substates required for allosteric function and/or binding. Structural mapping of communities across different states indicated that F592 could anchor the central inter-protomer communities and link them to G614 site. For more flexible 1 RBD-up conformational states of S-G614, we observed loosening of these communities around F592, suggesting that this position could effectively alter communications in the S-G614 trimer structures (Figure 10). At the same time, we observed some reorganization of the interfacial communities in the SD1-S2 region that harbors most of the hinge positions (Figure 10C,D). These network changes reflect strengthening of the interactions in the S2 subunit, while considerable mobility of the S1 regions can induce a reorganization of local communities. The results indicated that the regulatory control points could link together different local communities to form allosteric communication paths in the system. Hence, effective allosteric communications in functional states of the S-G614 trimer can be mediated by clusters of high centrality mediating residues. The network analysis revealed that the high centrality centers could be surrounded by clusters of neighboring residues featuring also fairly high betweenness values. As a result, functional sites critical for signal transmission between site of mutation and RBD regions could be shielded by neighboring residues that have sufficient communication capabilities to ensure broad communication routes and resilience in the fluctuating protein environment. Although mutations of functional residues may often result in a loss of activity, some of these changes could be rescued by neighboring residues that assume additional functional responsibilities in the altered interaction network.^156^

**Figure 8.**
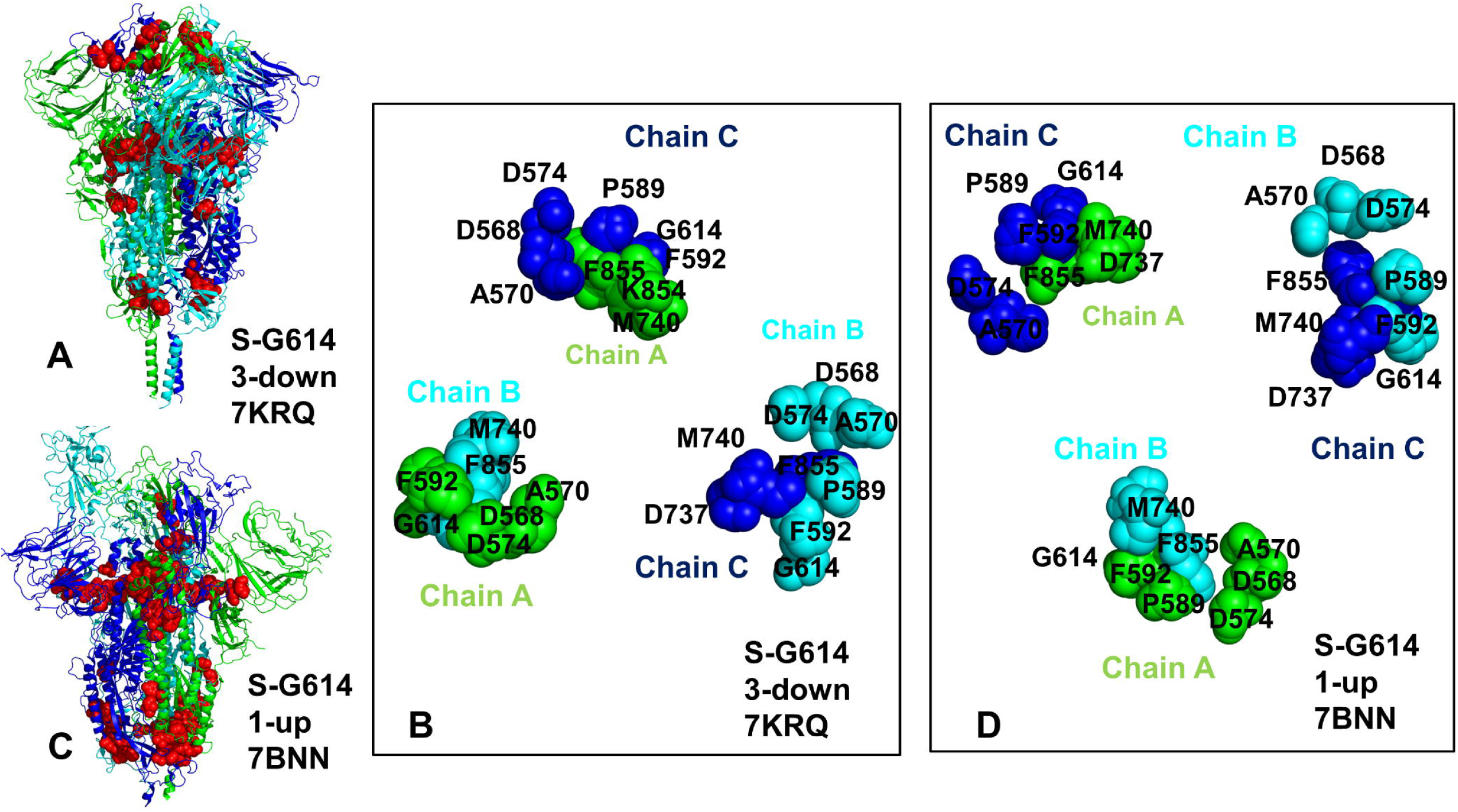
Network community analysis of the SARS-CoV-2 S-G614 closed and open states. Structural mapping of key local communities (residues comprising these modules are in red spheres) in the structure of SARS-CoV-2 S-G614 in the closed state, pdb id 7KRQ (A), and the structure 1 RBD-up open form, pdb id 7BNN (C). The SARS-CoV-2 S mutant protomer is shown in ribbons depicting the “down protomer” for the closed state on panel (A) and “up protomer” for the partially open state on panel (C). The S-G614 structures are shown in ribbons with the chain A in green, chain B in cyan, and chain C in blue colors. (B) A close-up of the inter-protomer hinge communities formed in the SARS-CoV-2 S-G614 in the closed state, pdb id 7KRQ. (D) A close-up of the inter-protomer hinge communities formed in the SARS-CoV-2 S-G614 in the 1 RBD-up open state, pdb id 7BNN. The communities are shown in spheres colored according to the respective chain. The residues that comprise the local communities are annotated. Note a partial fragmentation of local communities in the open form.

## Conclusions

The results of this study demonstrated that the D614G mutation may induce a considerable conformational adaptability of the open states without compromising folding stability and integrity of the spike protein. Using the local frustration profiling of the S-G614 conformational ensembles we found that conformational plasticity of the open S conformations can allow for moderate thermal rearrangements of the inter-protomer interactions and favor neutral frustration level near the mutational site and sites of circulating variants. We proposed a mechanism in which suboptimal inter-protomer interactions displaying specific frustration patterns may be exploited to control binding with the ACE2 and multiple binding partners. Functional dynamics analysis suggested that G614 site may serve as a versatile regulatory switch point that links or decouples the network of hinge clusters. The ensemble-based mutational energetic scanning of protein stability for the S-G614 substates suggests a significant mutational plasticity of the open S-G614 trimer. This study suggests that modulation of the energetic frustration at the inter-protomer interfaces by D614G mutation can serve as a mechanism for allosteric regulation in which the dynamic couplings between the site of mutation and the inter-protomer hinge of functional motions would modulate the inter-domain interactions, global changes in mobility and the increased stability of the open form. The network analysis showed that by promoting the enhanced mobility of the open states the D614G variant can induce allosteric communication routes that are robust to perturbations in the dynamic protein environment. The results of this study suggest that conformational plasticity and frustration-driven allostery of the S-614 protein are important functional mechanisms of this highly transmissible spike variant.

## DATA AND SOFTWARE AVAILABILITY

Crystal structures were obtained and downloaded from the Protein Data Bank (http://www.rcsb.org), accession numbers 7KRQ, 7BNM (the cryo-EM structures of SARS-CoV-2 S-G614 in the closed state), 7KRQ (the S-G614 intermediate state), 7KRR, 7BNN, 7EAZ, 7EB0, 7EB3 (1 RBD-up open states), and 7BNO, 7EB4, 7EB5 (2 RBD-up open states). All simulations were performed using NAMD 2.13 package that was obtained from website https://www.ks.uiuc.edu/Development/Download/. All simulations were performed using the all-atom additive CHARMM36 protein force field that can be obtained from http://mackerell.umaryland.edu/charmm_ff.shtml. The residue interaction network files in xml format were obtained for all structures using the Residue Interaction Network Generator (RING) program RING v2.0.1 freely available at http://old.protein.bio.unipd.it/ring/. The computations of network parameters were conducted using NAPS program available at https://bioinf.iiit.ac.in/NAPS/index.php and Cytoscape 3.8.2 environment available that can be downloaded from https://cytoscape.org/download.html. The rendering of SARS-CoV-2 S structures was done using the interactive visualization program UCSF ChimeraX package (https://www.rbvi.ucsf.edu/chimerax/) and Pymol (https://pymol.org/2/) . The scripts and programs for computational analysis are available in the GitHub site https://github.com/stevenagajanian/KInGAN.

## SUPPORTING INFORMATION

Figure S1 illustrates the domain organization for the full-length SARS-CoV-2 S protein. Figure S2 describes Functional dynamics of the SARS-CoV-2 S-D614 and S-G614 trimer structures in the locked closed, intermediate and open forms obtained using PCA of atomistic trajectories. Figure S3 shows the mutational sensitivity analysis of the G614 residue in the SARS-CoV-2 S-G614 closed state, intermediate state, and 1 RBD-up open form. Figure S4 details the mutational sensitivity analysis of the G614 residue in the SARS-CoV-2 S-G614 closed state, the 1 RBD-up state and 2 RBD-up open form. Figure S5 presents specifics of the mutational sensitivity analysis of the G614 residue in the ensemble of SARS-CoV-2 S-G614 open sub states. This material is available free of charge via the Internet at http://pubs.acs.org.

## AUTHOR INFORMATION

The authors declare no competing financial interest.

## Acknowledgment

The author acknowledges support by the Kay Family Foundation Grant A20-0032.

## ABBREVIATIONS

SARS: Severe Acute Respiratory Syndrome
RBD: Receptor Binding Domain
ACE2: Angiotensin-Converting Enzyme 2 (ACE2)
NTD: N-terminal domain
RBD: receptor-binding domain
CTD1: C-terminal domain 1
CTD2: C-terminal domain 2
FP: fusion peptide
FPPR: fusion peptide proximal region
HR1: heptad repeat 1
CH: central helix region
CD: connector domain
HR2: heptad repeat 2
TM: transmembrane anchor
CT: cytoplasmic tail

## For Table of Contents Use Only

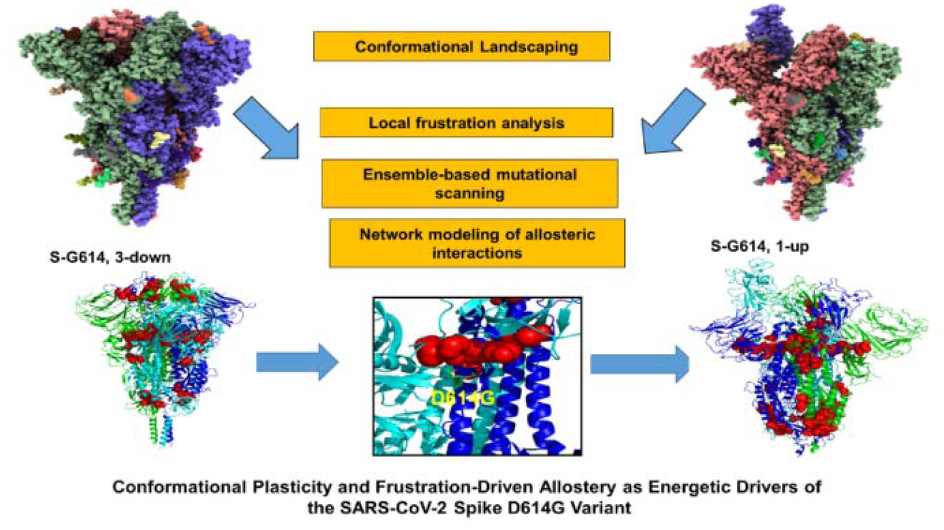

