## Supporting Figures S1-S5 for "The Landscape-Based Protein Stability Analysis and Network Modeling of Multiple Conformational States of the SARS-CoV-2 Spike D614 Mutant: Conformational Plasticity and Frustration-Driven Allostery as Energetic Drivers of Highly Transmissible Spike Variant"

### Supporting Information

### Atomistic Simulations and Landscape-Based Stability Analysis of Multiple Conformational States of the SARS-CoV-2 Spike D614G Variant: Mutation-Induced Conformational Plasticity of the Open State as Energetic Driver of Spike Functions

Gennady Verkhivker,[1,2]* Steve Agajanian[1], Ryan Kassab[1], Keerthi Krishnan[1]

[1] Keck Center for Science and Engineering, Graduate Program in Computational and Data Sciences, Schmid College of Science and Technology, Chapman University, Orange, CA 92866, United States of America

[2] Department of Biomedical and Pharmaceutical Sciences, Chapman University School of Pharmacy, Irvine, CA 92618, United States of America

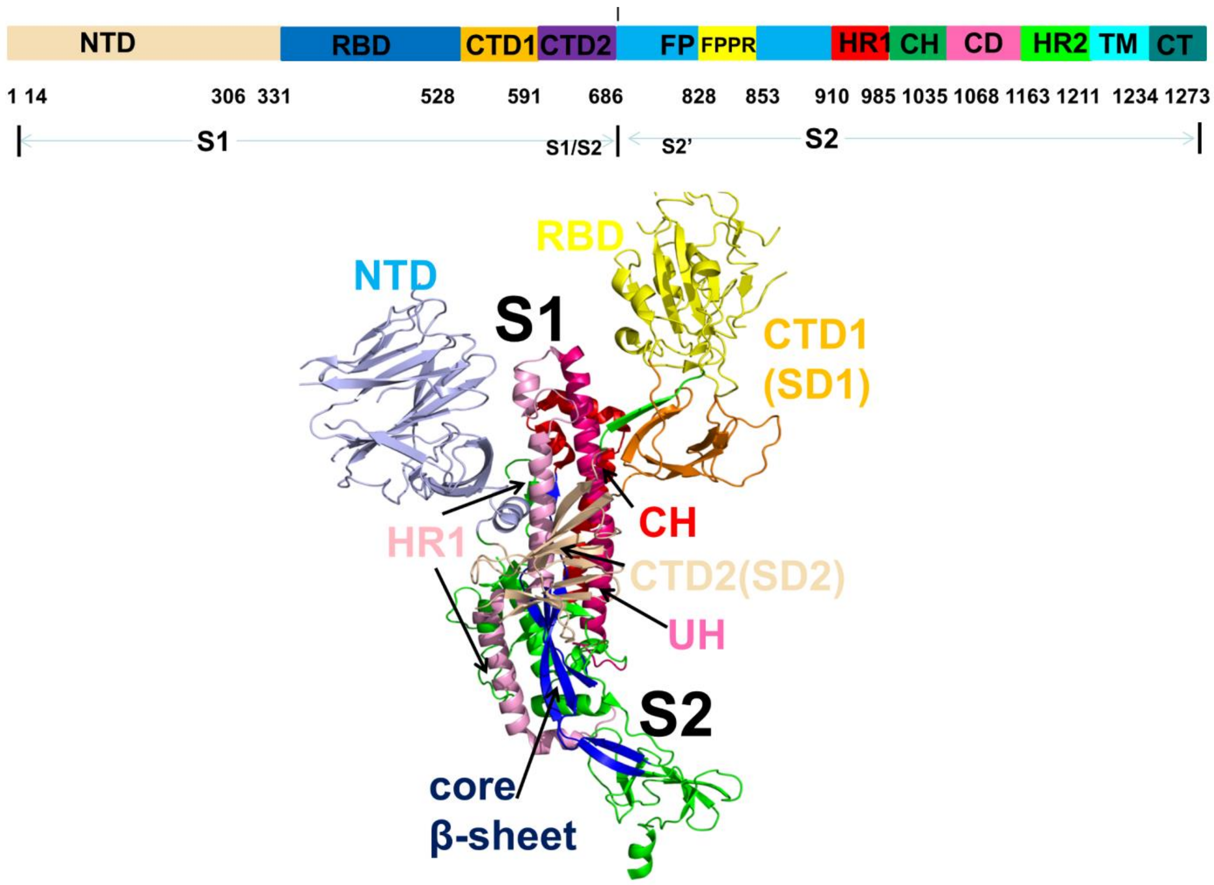


**Figure S1.** The domain organization for the full-length SARS-CoV-2 S protein. (A) The subunits S1 and S2 include NTD RBD, C-terminal domain 1(CTD1), C-terminal domain 2 (CTD2), fusion peptide (FP), fusion peptide proximal region (FPPR), heptad repeat 1 (HR1), central helix region (CH), connector domain (CD), heptad repeat 2 (HR2), transmembrane domain (TM), and cytoplasmic tail (CT). The subunits S1 regions : NTD (14-306) in light blue; RBD (331-528) in yellow; CTD1 ( 528-591) in orange; CTD2 (592-686) in wheat color ; upstream helix (UH) ( 736-781) in red; HR1 (910-985) in pink; CH (986-1035) in hot pink; core β-sheet (711-736, 1045-1076) (in blue).

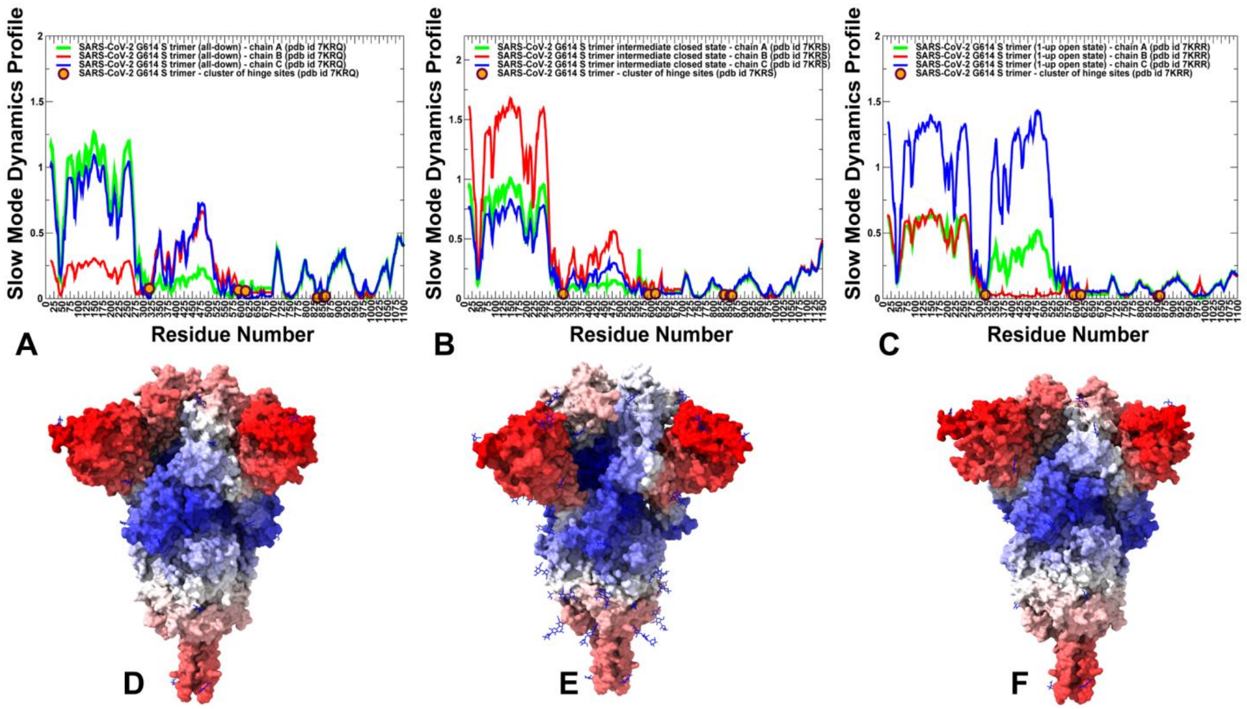


**Figure S2.** Functional dynamics of the SARS-CoV-2 S-D614 and S-G614 trimer structures in the locked closed, intermediate and open forms obtained using PCA of atomistic trajectories. The slow mode mobility profiles were averaged over the first three major low frequency modes. (A) The slow mode profile for the SARS-CoV-2 S-G614 trimer structure in the closed form (pdb id 7KRQ). (B) The slow mode profile for the SARS-CoV-2 S-G614 trimer in the intermediate closed form (pdb id 7KRS). (C) The slow mode profile for the SARS-CoV-2 S-G614 trimer structure in the 1 RBD-up open form (pdb id 7KRS). The profiles for protomer chains A, B and C are shown in green, red and blue lines, respectively. The positions of the hinge sites forming the inter-protomer cluster F318, F592, D614/G614, Y855, I856, and T859 are shown along the profiles in filled orange-colored circles. (D-F) Structural maps of the slow mode mobility profiles for these SARS-CoV-2 S-G614 structures. The position of mutational site D614G is highlighted in spheres and annotated. The structures are in sphere-based representation rendered using UCSF ChimeraX [90] with the rigidity-to-flexibility sliding scale colored from blue to red.

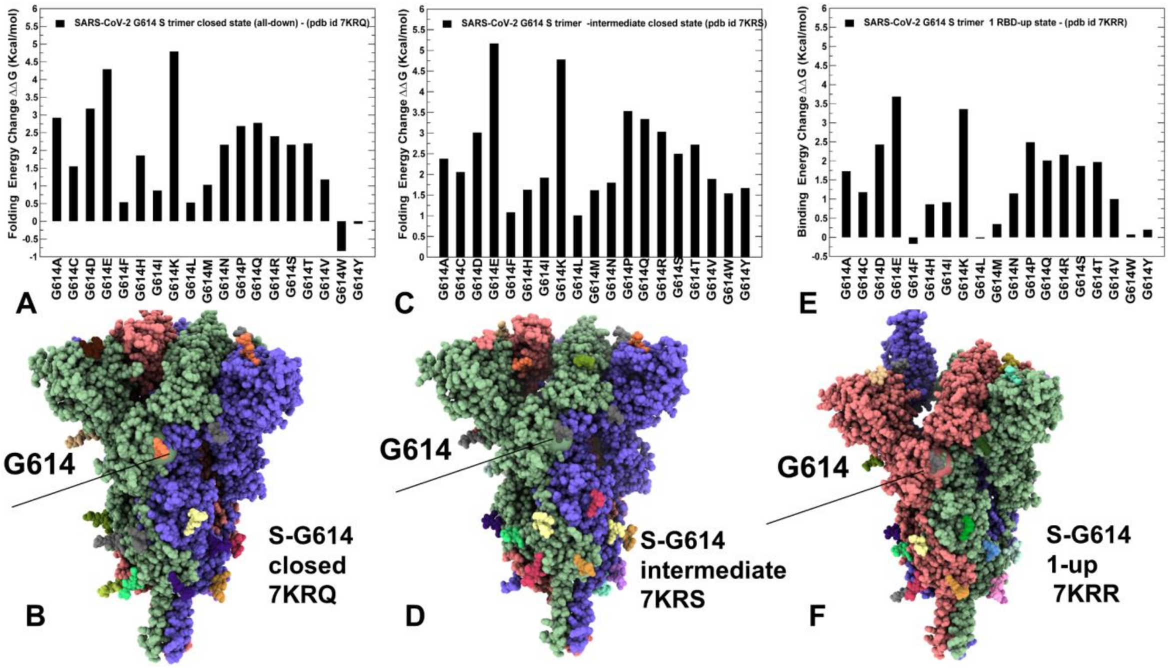


**Figure S3.** The mutational sensitivity analysis of the G614 residue in the SARS-CoV-2 S-G614 closed state, pdb id 7KRQ (A,B), in the intermediate state, pdb id 7KRS (C,D) and 1 RBD-up open form, pdb id 7KRR (E,F). The structures are shown in full spheres and colored with protomers A,B,C are colored in green, red and blue. The position of G614 is shown in spheres. The rendering of SARS-CoV-2 S structures was done using the interactive visualization program UCSF ChimeraX [90].

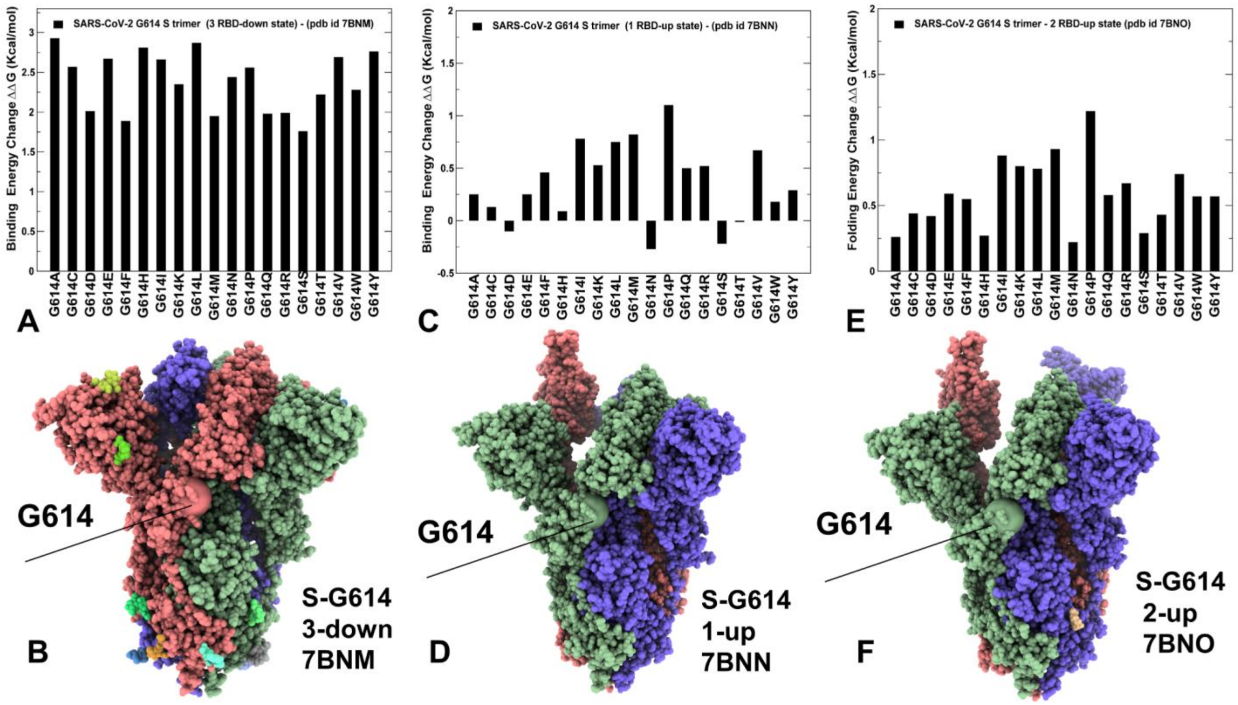


**Figure S4.** The mutational sensitivity analysis of the G614 residue in the SARS-CoV-2 S-G614 closed state, pdb id 7BNM (A,B), in the 1 RBD-up state, pdb id 7BNN (C,D) and 2 RBD-up open form, pdb id 7BNO (E,F). The structures are shown in full spheres and colored with protomers A,B,C are colored in green, red and blue. The position of G614 is shown in spheres. The rendering of SARS-CoV-2 S structures was done using the interactive visualization program UCSF ChimeraX [90].

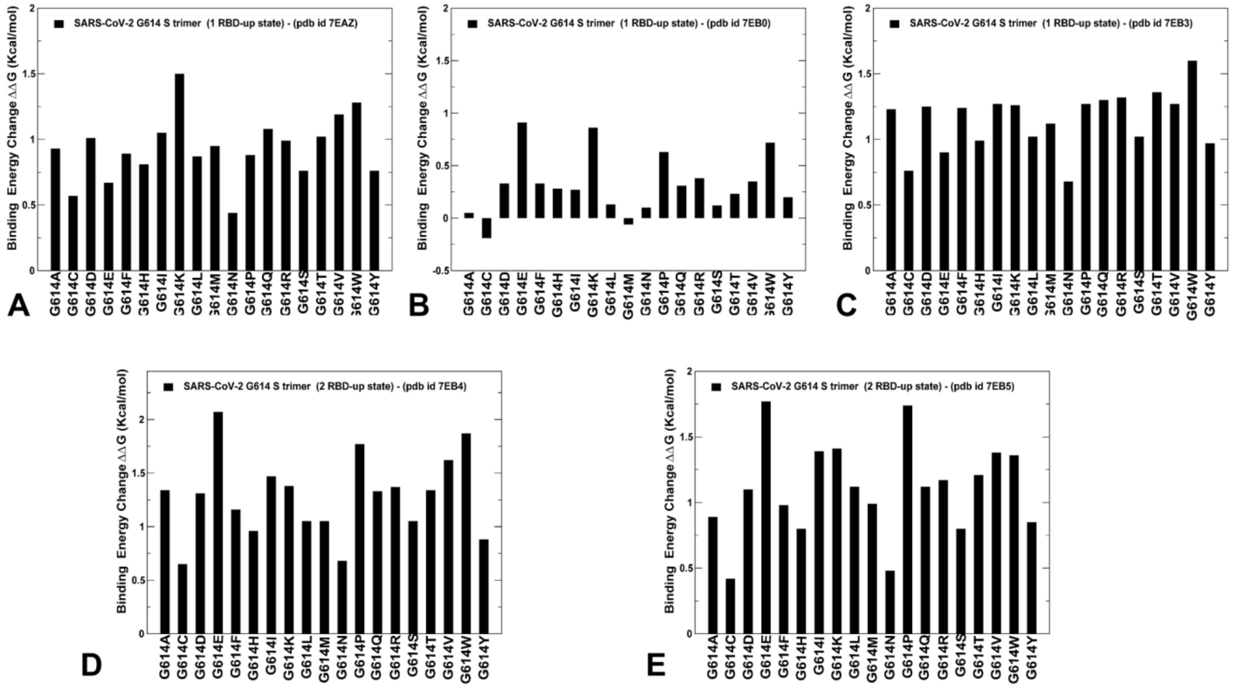


**Figure S5.** The mutational sensitivity analysis of the G614 residue in the SARS-CoV-2 S-G614 1 RBD-up open states (pdb id 7EAZ (A), pdb id 7EB0 (B), pdb OD 7EB3 (C)) and 2 RBD-up states (pdb id 7EB4 (D) and pdb id 7EB5 (E)).
